# Structural and Stereochemical Elucidation of Cyanochelin C, a Siderophore Associated with Novel Class of Cyanobacterial Acyl Hydrolases

**DOI:** 10.64898/2026.07.24.740629

**Authors:** Viviana Di Matteo, Lenka Marešová Štenclová, Berness P. Falcao, Pavel Hrouzek, Petra Urajová, Jan Mareš, Germana Esposito, Alfonso Mangoni, Valeria Costantino, Tomáš Galica

## Abstract

Iron is a key micronutrient that constrains microbial growth and productivity in many aquatic and terrestrial environments due to its limited bioavailability. Microorganisms evolved sophisticated acquisition strategies, including the production of siderophores, high-affinity iron-chelating molecules that facilitate iron solubilisation and uptake. Cyanobacteria, photosynthetic prokaryotes and major contributors to global primary production, also depend on iron as a cofactor to their core metabolic enzymes. However, very few cyanobacterial siderophores were described so far, and cyanobacteria remain an underxplored source of possibly novel siderophores. Here we report a novel cyanobacterial siderophore, cyanochelin C, that employs two β-hydroxyaspartate (β-OH-Asp) residues for iron chelation. We provide extensive nuclear magnetic resonance (NMR) and mass spectrometry (MS) evidence on the molecular structure and identify the corresponding biosynthetic gene cluster (BGC). Bioinformatic analysis of the BGC further revealed the presence of an acylase CcsQ clustering with a broader cyanobacteria-specific family of acylases associated with predicted siderophore-encoding BGCs. Discovery of cyanochelin C and its deacylation by CcsQ expands the known structural diversity of cyanobacterial siderophores and improves the understanding of important enzymatic reactions.

## Introduction

Cyanobacteria are photoautotrophic prokaryotes whose growth and reproduction depend on the availability of iron, an essential micronutrient required for numerous metabolic processes. Due to its specific chemical properties, iron is often not easily available to microorganisms. In oxic conditions, especially in marine environments, iron forms insoluble precipitates that sediment and leave open waters iron deprived. On the other hand, in arid soils, the nutrients are easily drained and in absence of water they cannot mix enough to cycle sufficiently. Another challenging and likely iron-deprived habitat represents bare rocks or man-made structures, which are often colonised by pioneering filamentous cyanobacteria.

Microbes evolved multiple mechanisms to overcome these difficulties and ensure sufficient supply of iron. One such mechanism is the secretion of siderophores - iron-chelating compounds that dissolve iron precipitates and scavenge trace iron from the environment. Cyanobacteria are frequently found in localities expected to suffer from long term or periodic iron deprivation, such as arid soils, moist bare rock surfaces, or pelagial of large water bodies. Genomic screenings for bioprospecting studies of secondary metabolites revealed numerous biosynthetic gene clusters that could possibly encode the production of complex high-affinity peptide siderophores. Yet the number of characterised cyanobacterial siderophores remains low. The currently known repertoire of cyanobacterial siderophores comprises 24 distinct structures: schizokinen, six variants of synechobactin, three variants of anachelins, six variants of cyanochelins, three leptochelins (5, 11, 12) and recently reported five variants of lusichelins (13).

Often, the siderophores share a common core structure and variants differ in minor structural features, especially in the length of the attached acyl chain or, possibly, its absence. Acylation is a frequent modification of cyanobacterial secondary metabolites and is thought to either limit the diffusibility of the compounds or to alter surface properties of the immediate environment (Galica et al 2017, Fewer et al 2021). The extensively studied bacterial siderophores, pyoverdines, often found in representatives of Pseudomonas and synthesised as acylated compounds. Only during maturation is their prolonged hydrophobic acyl residue cleaved off by an acyl hydrolase PvdQ (Drake and Gulick 2011). Marine bacteria of the genus Marinobacter often produce a suite of siderophores with various acyl residues at the N-terminus (Gauglitz et al 2014). Interestingly, in later stages of Marinobacter sp. DS40M6 batch culture, the acylated marinobactins (A-F) are converted to non-acylated marinobactin HG (Gauglitz et al 2014). Subsequent study identified BtnA (AJD87516.1) and MhtA (ENO13542.1) acylases responsible for the deacylation of marinobactins (Kem et al 2015, Kem and Butler 2015). While pyoverdines require deacylation by PvdQ for proper function and the gene encoding it is included in pyoverdine BGCs (Drake and Gulick 2011), the acylated marinobactins A-E are considered functional and their deacylation seems optional (Gauglitz et al., 2014). Both described marinobactin acylases, BtnA and MhtA, are not considered as part of the marinobactin BGC and their primary substrate may be something else. Apart from marinobactins, MhtA can deacylate C12 acyl-homoserine lactone (AHL), and so can PvdQ (Kem et al 2015, Bokhove et al 2009). Degradation of AHLs interferes with quorum sensing of various bacteria and is also known as quorum quenching. While quorum sensing via AHLs has not been described in cyanobacteria, Romero and colleagues have identified a functional AHL acylase, designated as AiiC, in Anabaena (Nostoc) sp. PCC 7120 that can interfere with quorum sensing of co-habiting bacteria (Romero et al 2008).

In our previous study, we identified and characterised the acylated β-OH-Asp-containing cyanobacterial siderophores cyanochelins A and B, as well as several minor structural variants (Galica et al., 2021; Falcão et al., 2025). Ever since, our systematic effort to elucidate the diversity of cyanobacterial secondary metabolites has included a special focus on β-OH-Asp-siderophore-encoding BGCs combined with additional, less-explored tailoring enzymes. Given that many bacterial siderophores occur in both acylated and non-acylated forms, we sought to determine whether and how cyanobacteria produce non-acylated cyanochelins or related peptide siderophores. In the present, study we describe a siderophore-encoding BGC from soil-dwelling filamentous cyanobacteria *Myxacorys* spp. The BGC includes modules for incorporation of β-OH-Asp, as well as an unusual acylase that, as we show later, belongs to a novel family of cyanobacterial siderophore acylases. Most importantly, we report the isolation and full structural stereochemical elucidation of cyanochelin C. Using a powerful multidisciplinary strategy for natural product characterisation, by combining direct stereochemical information obtained through chemical analysis with biosynthetic insights derived from genomic data, we obtained complementary and mutually reinforcing evidence. This synergistic framework significantly enhances the accuracy, reliability, and overall robustness of stereochemical assignments.

## Results

### Biosynthetic gene cluster encoding production of cyanochelin C

Our recent genome mining efforts identified multiple strains with candidate cyanochelin-like biosynthetic gene clusters in representatives of Leptolyngbyaceae, including two strains of *Myxacorys* spp. - *M. chilensis* ATA2-1-KO14 and *M. californica* WJT36-NPBG1 (Pietrasiak et 2019). A putative siderophore-producing BGC was identified by AntiSMASH in both genomes (NCBI: JAHHHP010000001.1 and JAFJZQ010000037.1) with 89,6% nucleotide similarity across the whole cluster. The NRPS BGC consists of nine biosynthetic ORFs (Fig. 1) and an additional Ntn-CA acylase (NCBI: MBW4537642.1). With the notable exception of the acylase the genes for biosynthesis of cyanochelin C are organised in one direction similarly to other cyanochelin BGCs we have examined so far.

The first ORF (*ccsAXE,* naming reflects major cyanochelin A genes) includes a fatty acid-AMP ligase (FAAL) domain and 2 NRPS modules for incorporation of L-valine incorporation and cyclisation of L-threonine (*E*). The second ORF (*ccsF*) encodes an NRPS module for activation of lysine and its subsequent epimerasion to D-lysine. The third gene (*ccsG*) encodes a β-alanine ( β-Ala) specific NRPS module and the fourth ORF (*ccsR*) encodes a transaminase to ensure the supply of β-Ala. ORFs 5-7 (*ccsH_1/2_*, *ccsH_2/2_* and *ccsI*) encode an NRPS module for incorporation of β-OH-Asp. ORF 8 (*ccsJK*) contains modules for Gly incorporation as well as a C-domain and TauD (taurine dioxigenase D) that together with ORF9 (*ccsL*) participate in biosynthesis of the last amino acid, a C-terminal β-OH-Asp or possibly 2,4-dihydroxy-3-aminobutyric acid (DHABA). The putative product is released by the thiodiesterase domain at ORF9.

The gene encoding the final NRPS module (ORF9/*ccsL*) is followed by a set of oppositely oriented genes encoding glycosyl transferases and exopolysacharide biosynthesis enzymes, for which we found no evidence of involvement in cyanochelin C biosynthesis. At the opposite end of the BGC, upstream of the core biosynthetic genes and transcribed in the opposite orientation, lies a gene encoding an acylase (NCBI: MBW4537642.1, *ccsQ).* Adjacent, to *ccsQ* there is a complete siderophore transporter cassette (Fig. 1., Table S1), providing additional evidence that the product of these BGCs functions as a siderophore. The transporter genes were annotated according to their putative functional counter-parts in the proposed cyanochelin B transporter cassette. With the exception of cctG and cctH, which are likely involved in active export, all transporter components could be confidently assigned. The transporter cassette is flanked on its outer side by genes encoding homologues of the iron exporters FetA and FetB. In E*scherichia* coli, the FetAB system was demonstrated to facilitate oxidative stress relief by exporting iron ions from the cell (Nicolau et al 2013, ctg1_3: FetA, ctg1_2 FetB).

**Fig. 1.**
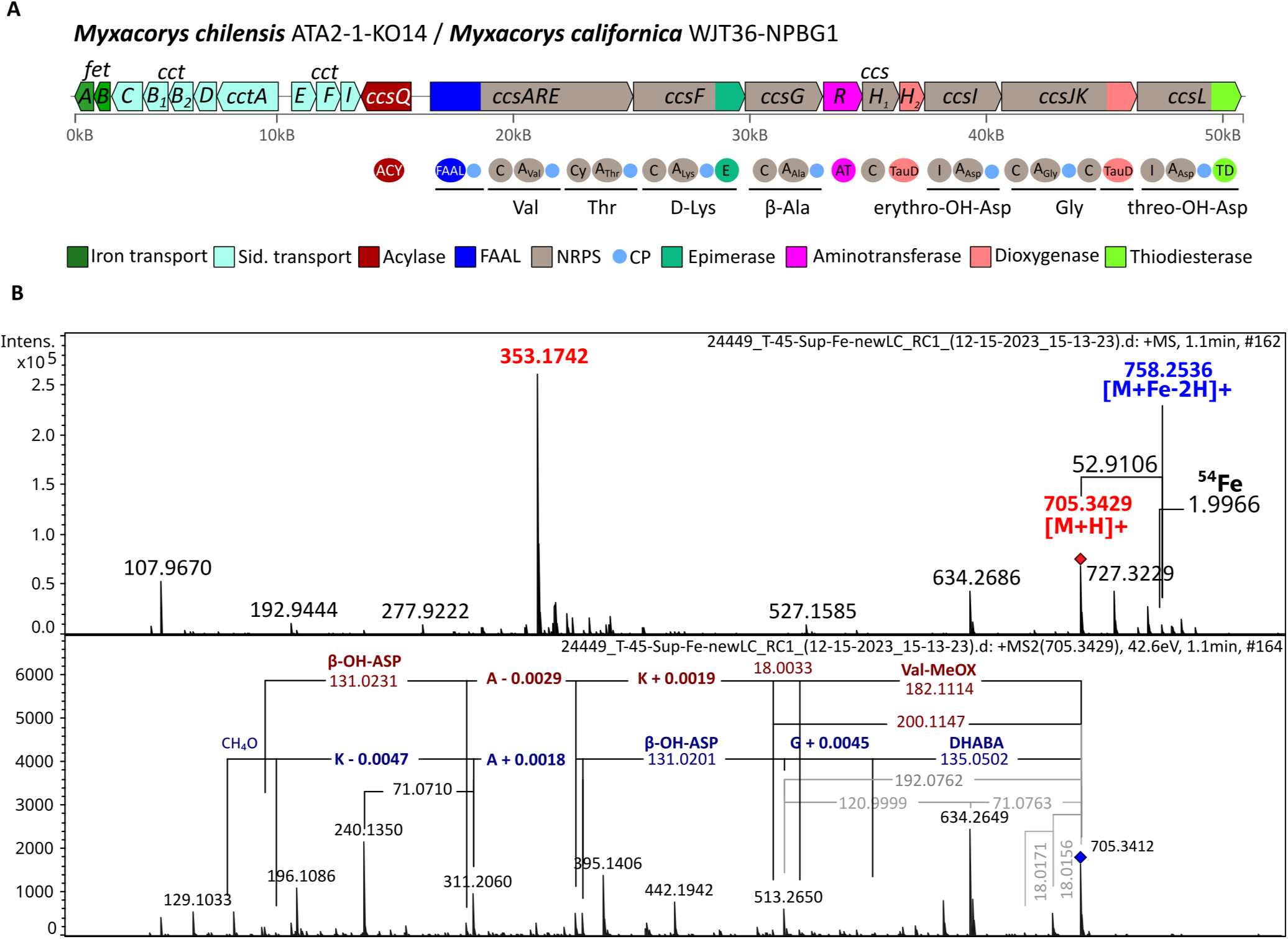
Interpretation of the cyanochelin C biosynthetic gene cluster (BGC) and identification of its product. (**A**) Organisation of the cyanochelin C BGC. The cluster comprises nine open reading frames (ORF) encoding an non-ribosomal peptide synthetase (NRPS) assembly line and and the maturation acylases CcsQ, located upstream of the core NRPS genes in opposite orientation. A transporter cassette, likely involved in siderophore import and export, is located just next to the BGC. Adjacent *fetAB* genes, involved in iron transport, are highlighted in green. Gene names follow the nomenclature established for cyanochelin A BGC to maintain consistency across cyanochelin variants. (**B**) HPLC-HRMS analysis of the iron-starved M*yxacorys* chilensis. Cyanochelin C and its iron complexes were detected in MS1 spectra, while MS/MS fragmentation analysis supported the peptide sequence predicted from the BGC.

### Identification and isolation of Cyanochelin C

Pietrasiak and colleagues reported the isolation of several *Myxacorys* strains, including *M. chilensis* ATA2-1-KO12 and ATA2-1-KO14 from the same environmental sample collected in the Atacama Desert in 2009 (Pietrasiak et al 2019). Strain ATA2-1-KO14 was subsequently designated as the type (reference) strain and its genome was sequenced (Ward et al., 2021). Given their common origin and high morphological similarity, the two strains are likely clonal, or differ minimally (Pietrasiak et al 2019). To induce iron stress and stimulate siderophore production, both strains were cultivated in iron-depleted BG-11 medium. Siderophore production was assessed in biomass and culture supernatants using the Chrome-azurol-S assay (CAS, Schwyn and Neilands, 1987). The CAS-positive cultures were analysed by HPLC/HRMS. To facilitate siderophore identification, FeCl_3_ (5µM) was added to an aliquot of the extract and analysed separately. Compounds (features) were detected using mzMine 3.9 and screened by a custom Python script to identify features that: 1) exhibit an isotopic pattern indicating the presence of bound iron (isotope Fe^54^ 6.5% of Fe^56^) and 2) have a matching feature without the bound iron. Several of the best scoring candidates were analysed manually, yielding the candidate compound with a siderophore at *m/z* 705.3429 and an iron-siderophore complex at *m/z* 758.2536. MS^2^ spectra obtained at 42eV provided a series of fragments corresponding to neutral losses of predicted amino acids from each side of the linear NRPS peptide and indicated the presence of β-OH-Asp. The compound was found in both strains of *Myxacorys* with higher yields from ATA2-1-KO12, which was chosen as material for compound isolation.

To isolate the compound, spent medium (3 L) of iron starved *Myxacorys chilensis* ATA2-1-KO12 was processed by one-step solid phase extraction (SPE) and the obtained extract was subjected to HPLC fractionation to yield 17.6 mg of the candidate compound.

### Structure elucidation of cyanochelin C

The purified compound was subjected to a comprehensive set of 1D and 2D homo- and heteronuclear NMR experiments, complemented by high-resolution mass spectrometry (HRMS) analysis. The structural elucidation of cyanochelin C was further supported by bioinformatic analysis of its putative biosynthetic gene cluster (BGC) as well as by Marfey’s and Murata’s methods for stereochemical determination. The HRMS spectrum of cyanochelin C (Fig.1) displayed a protonated molecular ion [M+H]+ at *m/z* 705.3429, consistent with the molecular formula C_28_H_49_N_8_O_13_. Tandem mass spectrometry (MS/MS) experiments employing collision-induced dissociation were performed to further investigate the peptide sequence. The MS/MS spectrum of the ion at *m/z* 705.3412 showed sequential losses of individual amino acid residues starting from the N-terminus, generating fragment ions at *m/z* 634.2649 (x6, loss of Val), 523.2367 (y5, loss of methyloxazoline–CO), 395.1406 (y4, loss of Lys), 324.1036 (y3, loss of β-Ala), and 193.0817 (y2, loss of β-OH-Asp). This fragmentation pattern was consistent with the substrate specificity and module organisation of the adenylation (A) domains identified in the NRPS assembly line.

The peptide substructure was further supported by the NMR data (Table. 1, Fig. S1-S8), which revealed five signals attributable to α-protons of amino acid residues (δ _H_ of H-3: 4.30 ppm, H-8: 4.62 ppm, H-15: 4.28 ppm, H-21: 4.39 ppm and H-25: 3.84 ppm). In addition, two methylene proton signals at δ_H_ 3.89 ppm (H-6a) and 3.85 ppm (H-6b) were consistent with a glycine residue. Signals corresponding to a β-Ala residue were also observed, including methylene resonances at δ_H_ 2.56 ppm (H-12a b), 3.43 ppm (H-13a), and 3.51 ppm (H-13b). Two oxygenated methine protons at C-2 and C-9 resonating at δ_H_ 4.12 ppm and 4.21 ppm, respectively, supported the presence of hydroxyl groups at the β-positions of Asp residues. Furthermore, the observation of methylene protons bound to a hydroxyl group (δ_H_ of H-4a: 3.58 ppm and H-4b: 3.68 ppm), together with COSY and HMBC correlations, indicated that the C-terminal residue corresponds to 2,4-dihydroxy-3-aminobutyric acid (DHABA). This residue is likely derived from a β-OH-Asp unit that undergoes reduction to a primary alcohol via a reductive thioesterase domain encoded within the Asp-specific NRPS module, located in the final gene of the BGC (Fig. 1). Moreover, HMBC correlations supported the presence of a methyl-oxazoline ring (MeOx) formed by cyclisation of the threonine side chain with the carbonyl group of the valine residue. Specifically, H-21 (δ_H_ 4.39 ppm) exhibited an HMBC correlation with the valine carbonyl carbon C-25, while H-22 (δ_H_ 4.85 ppm) from the threonine side chain correlated with the valine α-carbon C-24. The presence of a methyl-oxazoline moiety is also consistent with the biosynthetic prediction, as the NRPS assembly line contains a threonine-specific module bearing a cyclisation (Cy) domain, which typically catalyses the formation of oxazoline heterocycles.

To enable detection of amide NH resonances, additional NMR spectra were recorded in H_2_O/D_2_O (9:1) containing 20 mM phosphate buffer, as cyanochelin C was soluble exclusively in aqueous media (Table. 1).

**Table. 1.** NMR data were recorded at 700 MHz in D_2_O at 398K. Chemical shifts (δ_H_) values of amide NH protons were instead obtained from spectra acquired in H_2_O/D_2_O (9:1) containing 20 mM phosphate buffer. Cyanochelin C thus consists of [Val^1^, MeOx^2^, Lys^3^, β-Ala^4^, β-OH-Asp^5^, Gly^6^, DHABA^7^] and contains 9 stereocenters.

| Residue | Position | | $\delta_H$ [mult., J (Hz)] | $\delta_C$ [mult.] | HMBC ($^1H \rightarrow ^{13}C$) |
| --- | --- | --- | --- | --- | --- |
| DHABA | 1 |  |  | 178.2, C |  |
|  | 2 |  | 4.12, d (2.5) | 70.6, CH | 1, 3, 4 |
|  | OH-2 |  |  |  |  |
|  | 3 |  | 4.30, ddd (8.2, 7.4, 2.5) | 53.8, CH | 1, 2, 4, 5 |
|  | NH-3<sub>(H<sub>2</sub>O)</sub> |  | 7.52, d (9.4) |  |  |
|  | 4 | a | 3.58, dd (11.4, 6.4) | 61.1, CH<sub>2</sub> | 2,3 |
|  |  | b | 3.68, dd (11.6, 7.7) |  |  |
| Gly | OH-4 |  |  |  |  |
|  | 5 |  |  | 171.1, C |  |
|  | 6 | a | 3.89, d (17.3) | 42.5, CH | 5,7 |
|  |  | b | 3.95, d (17.3) |  | 5,7 |
|  | NH-6<sub>(H<sub>2</sub>O)</sub> |  | 8.25, m |  |  |
| L-β-OH-Asp | 7 | | | 171.3, C | |
|  | 8 |  | 4.62, d (5.2) | 57.0, CH | 7, 9, 10, 11 |
|  | 9 |  | 4.21, d (5.2) | 72.2, CH | 7, 8, 10 |
|  | OH-9 |  |  |  |  |
|  | 10 |  |  | 176.7, C |  |
|  | NH-8<sub>(H<sub>2</sub>O)</sub> |  | 8.13, d (5.9) |  |  |
| β-Ala | 11 | | | 174.0, C | |
|  | 12 | a | 2.56, t (6.3) | 34.5, CH<sub>2</sub> | 11, 13 |
|  |  | b | 2.56, t (6.3) |  | 11, 13 |
|  | 13 | a | 3.43, m | 35.5, CH<sub>2</sub> | 11, 12, 14 |
|  |  | b | 3.51, m |  | 11, 12, 14 |
|  | NH-13<sub>(H<sub>2</sub>O)</sub> |  | 8.21, t (11.6, 5.6) |  |  |
| D-Lys | 14 |  |  | 173.3, C |  |
|  | 15 |  | 4.28, dd (8.2, 6.4) | 53.5, CH | 14, 16, 17, 20 |
|  | 16 | a | 1.74, m | 30.6, CH<sub>2</sub> | 14, 15, 17, 18 |
|  |  | b | 1.77, m |  | 14, 15, 17, 18 |
|  | 17 | a | 1.37, m | 21.9, CH<sub>2</sub> | 15, 16, 18, 19 |
|  |  | b | 1.41, m |  | 15, 16, 18, 19 |
|  | 18 | a | 1.66, quint (15.4, 7.7) | 26.0, CH<sub>2</sub> | 16, 17, 19 |
|  |  | b | 1.66, quint (15.4, 7.7) |  | 16, 17, 19 |
|  | 19 | a | 2.97, t (7.7) | 38.8, CH<sub>2</sub> | 17, 18 |
|  |  | b | 2.97, t (7.7) |  | 17, 18 |
|  | NH-15<sub>(H<sub>2</sub>O)</sub> |  | 8.13, d (5.9) |  |  |
|  | NH<sub>2</sub>-19 |  |  |  |  |
| L-Thr | 20 |  |  | 172.6, C |  |
|  | 21 |  | 4.39, d (6.9) | 73.0, CH | 20, 22, 23, 24, 25 |
|  | 22 |  | 4.85, quint (13.0, 6.5) | 81.6, CH | 20, 24 |
|  | 23 |  | 1.44, d (6.3) | 20.1, CH<sub>3</sub> | 21, 22 |
|  | 24 |  |  | 167.7, C |  |
| L-Val | 25 |  | 3.84, m | 54.4, CH |  |
|  | 26 |  | 2.19, m | 30.1, CH | 24, 25, 27, 28 |
|  | 27 |  | 1.00, d (6.9) | 17.9, CH<sub>3</sub> | 25, 26 |
|  | 28 |  | 1.02, d (6.9) | 17.0, CH<sub>3</sub> | 25, 26 |
|  | NH<sub>2</sub>-25 |  |  |  |  |

The absolute configuration of the amino acid residues was determined using the advanced Marfey’s method (Marfey 1984), a widely applied approach for stereochemical assignment based on chiral derivatisation followed by chromatographic analysis and critically corroborated by detailed analysis of the corresponding NRPS modules. The integration of chemical and genomic data provided complementary and mutually reinforcing evidence, thereby ensuring a robust and reliable stereochemical characterisation. In this way, the presence of L-Val (*S*-Val), L-Thr (2*S*,3*R*-Thr), and D-Lys (*R*-Lys) was demonstrated (Fig. S9 and Fig. S10) The detection of D-Lys is consistent with the presence of an epimerase domain within the NRPS module responsible for incorporating the Lys3 residue, clearly indicating that stereochemical inversion occurs during its biosynthetic assembly.

The relative configurations of the β-OH-Asp and DHABA residues were initially unclear and were therefore investigated through detailed analysis of the NMR data (Matsumori et al. 1999). Examination of the DHABA moiety revealed that proton H-3, resonating as a doublet of doublets of doublets at δ _H_ 4.30 ppm, shows a ^3^*J*_H,H_ coupling constant of 2 *Hz* with H-2. Comparison of the multiplicities of H-3 in the ^1^H NMR spectrum and in the 1D slices of the HMBC spectrum recorded at δ_C_ 178.2 (C-1) indicated that the ^3^*J*_C,H_ coupling constant between H-3/C-1 is small. In addition, the magnitudes and sign of ^3^*J*_C,H_ coupling constant between H-2/C-4 were determined from the HSQC-HECADE spectrum and were likewise found to be small.

Furthermore, the magnitude and signs of ^2^*J*_C,H_ coupling constants between H-3/C-2 and H-2/C-3 were also measured from the HSQC-HECADE spectrum, and detected as small. Altogether, these data supported a *threo* relative configuration for the DHABA residue (Fig. 2).

A similar analysis was performed for the β-OH-Asp residue. Proton H-8, resonating as a doublet at δ_H_ 4.62 ppm, is coupled to H-9 with a ^3^*J*_H,H_ coupling constant of 5.2 *Hz*. Comparison of the multiplicities of H-8 and H-9 in the 1H NMR spectrum and in the 1D sections of the HMBC spectrum recorded at δ_C_ 171.3 (C-7) and δ_C_ 176.7 (C-10), respectively, showed a small ^3^*J*_C,H_ coupling constant between H-8 and C-10 and a medium coupling between H-9 and C-7 (Fig. 2). In addition, the magnitudes and signs of ^2^*J*_C,H_ coupling constants between H-9/C-8 (−4.1 Hz) and H-8/C-9 (−4.2 Hz) were recorded using HSQC-HECADE spectrum and were both medium. According to the empirical models proposed by Murata, a larger ^2^*J*_C,H_ value would typically be expected for the H-9/C-8 pair in the *erythro* configuration. However, in the present case, C-9 is bound to a nitrogen atom rather than to a hydroxyl group, as in the reference systems described by Murata. The lower electronegativity of nitrogen compared to oxygen may therefore influence the magnitude of the observed coupling constants. Nevertheless, considering that the observed ^3^*J* coupling pattern is consistent only with the *erythro* model reported by Murata, the β-OH-Asp residue was assigned an *erythro* relative configuration.

Bioinformatic analysis of the BGC supports these assignments. In particular, the NRPS module responsible for incorporating the β-OH-Asp residue lacks an epimerization domain, indicating retention of the L configuration. Accordingly, the stereochemistry of these residues was assigned as L**-**β-OH-Asp (*8S,9R*) and (*2R,3S*)-DHABA.

**Fig. 2.**
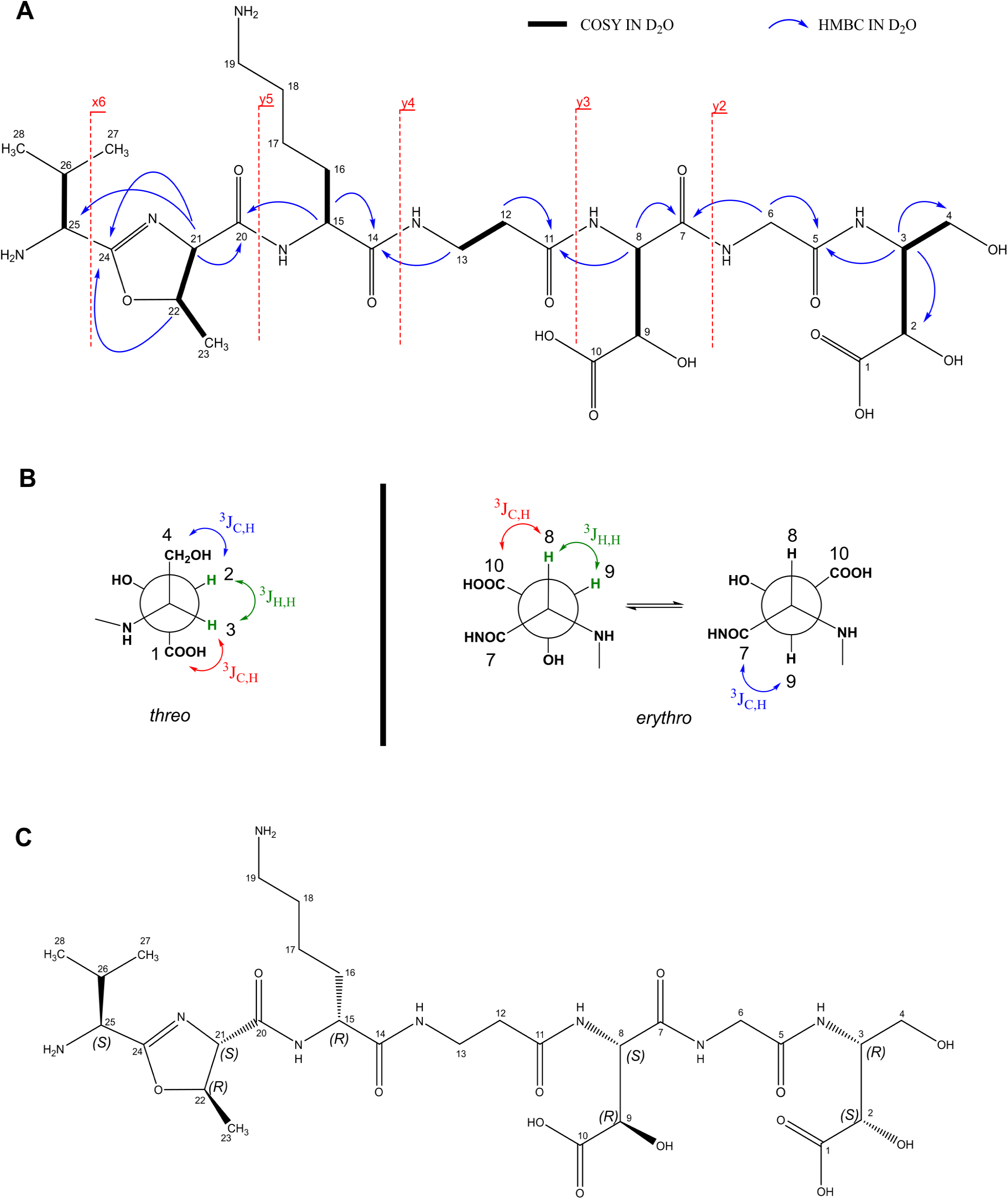
Cyanochelin C: **(A)** Key 2D NMR correlations (D_2_O) and diagnostic MS/MS fragments to establish the amino acid sequence and connectivity. **(B**) Newman projections showing the *threo* and *erythro* relative configurations of the DHABA and β-OH-Asp residues, with the ^3^*J*_H,H_ and ^3^*J*_C,H_ coupling constants used to discriminate between the two diastereomeric relationships. **(C)** Full stereostructure.

### Characterization of siderophore maturation acylase

The presence of FAAL within the BGC led to the initial prediction that the biosynthetic product would contain an N-terminal acyl moiety. However, the siderophores detected in the iron starved cultures did not include acyl residue. An in-depth inspection of the BGC, revealed the presence of a putative acylase (CcsQ) with significant sequence similarity to PvdQ acylase, known to be involved in maturation of pyoverdines. Phylogenetic analysis of CcsQ and N-terminal nucleophile acylases confirmed the homology of CcsQ to PvdQ and marinobactin acylases (BntA and MhtA/B) within a broader lineage of bacterial AHL acylases, including AiiC from Anabaena sp. PCC 7120 (Fig. 3). However, CcsQ exhibited an even closer evolutionary relationship to a set of structurally described cephalosporin/ glutaryl-7-ACA acylases (O86089, P07662, AAN39264, Q9L5D6). Taking advantage of accumulated structural data on similar acylases, tertiary structure of CcsQ was modelled using AlphaFold3 including a Leu-Ser-Lys-Ala-Asp-Ala-Asp and myristic acid as ligands, mimicking a situation just after acyl cleavage (Fig. 3). The modelling yielded high-confidence (pTM 0.96) folding of CcsQ and placed the myristic acid into a binding pocket (CcsQ-myristic acid pair ipTM 0.96) so that the carboxylic functional residue is oriented towards the active site (CssQ: Ser 161, His 183, Val 229 and Asn 403; PvdQ: Ser 217, His 239, Val 286 and Asn 485) and the first amino acid (Leu) of the modelled peptide (CcsQ-peptide ipTM 0.69). This configuration supports the hypothesis that the identified acylase is responsible for the release of the fatty acid from the nascent peptide.

**Fig. 3.**
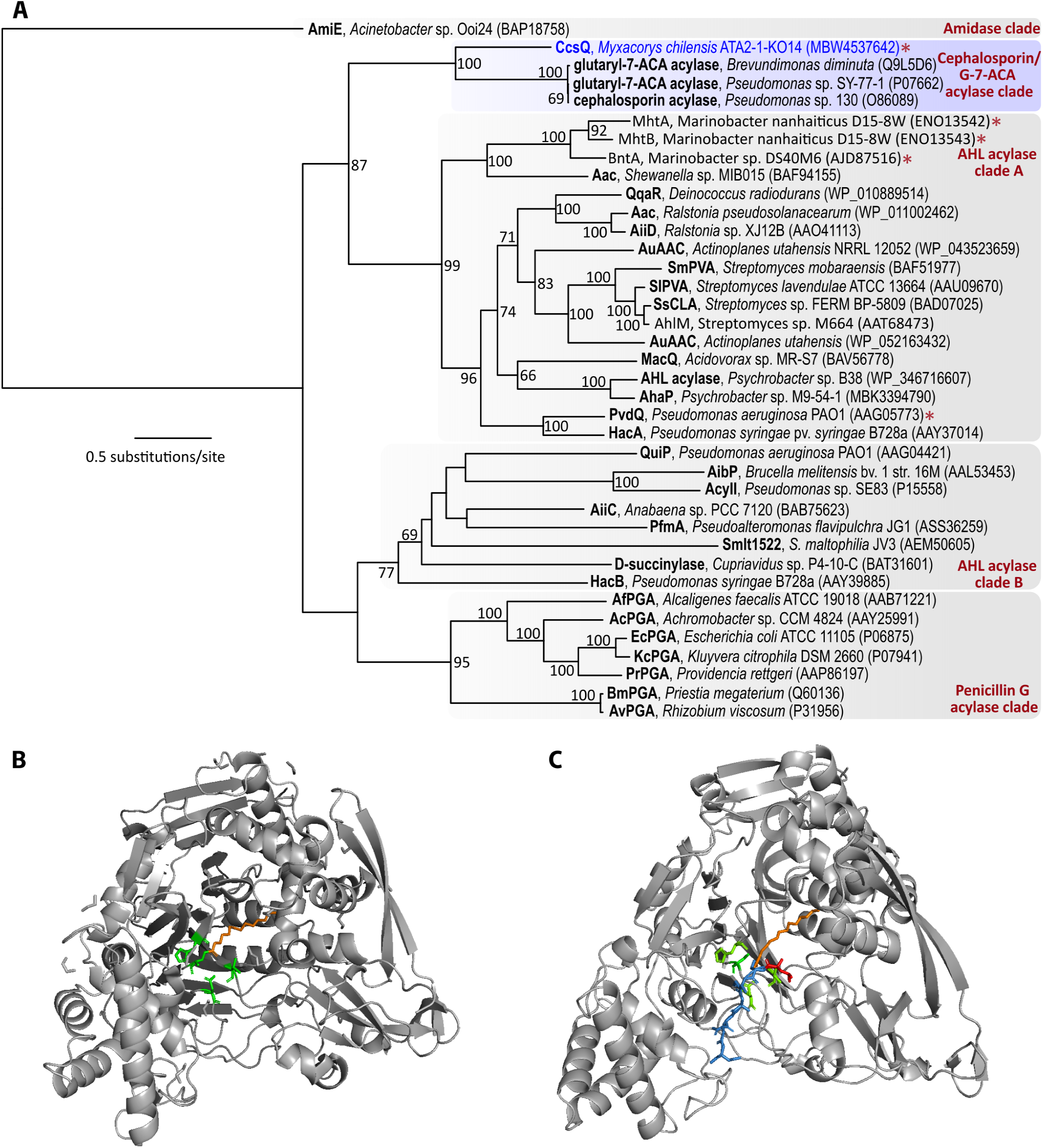
Phylogenetic and functional assessment of CssQ: **(A)** *Phylogenetic analysis of bacterial N-terminal nucleophile acylases -* Cyanochelin C acylase (CcsQ) was resolved as a distant homologue of several groups of AHL acylase and penicillin acylase-like hydrolases. The closest experimentally characterised enzymes are cephalosporin/ glutaryl-7-ACA acylases; the second-closest sister group of AHL acylase-like enzymes contains the pyoverdine acylase PvdQ and marinobactin acylases BntA and MhtA/B. The tree was calculated using the Maximum Likelihood method with 1000 bootstrap repetitions and rooted with AmiE (amidase), branch support values greater than 50 % are shown at the nodes. **(B)** Structure of PvdQ with myristoic acid (orange) bound to active site (green) Ser (Ser *β*217) as resolved by Drake and Gulick 2011 (3SRA). **(C)** CcsQ as modeled by Alphafold 3 (without 36 AA long N-terminal signal sequence) viewed from similar angle as PvdQ in **B.** In addition, peptide LSKADGD corresponding to unmodified peptide core of the siderophore is depicted in blue, except the N-terminal Leu, in red, that would be the residue connected to myristoic acid. The situation depicted mimics moments just after deacylation.

### Distribution of N-terminal nucleophile acyl hydrolases across the cyanobacterial phylum

To explore the broad phylogenetic landscape of the CchQ acylase within all bacteria, we executed BLAST search and general sequence-similarity network (SSN) analysis based on 997 retrieved closest BLAST hits. The results indicated that cyanobacterial homologues of CcsQ form a coherent group separated from homologous proteins in other bacterial phyla, with most abundant hits originating from Pseudomonadota, Bacteriodota, and Fidelibacterota, and few representatives from Gemmatimonadota, Acidobacteriota, and several other phyla (Fig. 4.).

We were interested to see if some cyanobacterial acylases could possibly be specialised for siderophore maturation. Inspection of our previously published dataset of cyanochelin-like BGCs (Galica et al 2021) revealed the presence of an acylase in 22 out of 31 siderophore BGCs. To expand our search, we used the 22 putative siderophore-associated acylases to construct a position specific scoring matrix (PSSM) and used it as a query for a subsequent PSI-BLAST search against NCBI’s database of clustered non-redundant proteins (ClusteredNR). Among the first 1 000 hits, 259 sequences were assigned to Cyanobacteria, with the remaining homologs distributed across bacterial taxa from phyla Pseudomonadota, Bacteriodota, Fidelibacterota, Gemmatimonadota and Acidobacteriota. Phylogenetic tree constructed from these results largely reflects the established SSN grouping according to enzyme function and phylogenetic origin (Fig. 4.). Notably, the closest homologs to cyanobacterial acylases retrieved by PSI-BLAST, in contrast to the naive SSN analysis, were uncharacterized acylases from Acidobacteriota, which partially interspersed within the few cyanobacterial representatives clustering at the basis of the lineage (Fig. 4.). For further detailed phylogenetic analysis we employed sequences of 259 cyanobacterial proteins and as an outgroup 31 closest CcsQ-homologues from Acidobacteriota (Fig. 5., Fig. S11). Multiple sequence alignments and data curation were performed to yield the final dataset of 292 taxa and 642 aligned amino acid positions that was subsequently used to construct the phylogenetic tree of N-terminal nucleophile hydrolases in cyanobacteria. Proteins included in the phylogenetic analysis were examined for their association to putative siderophore BGC. Genomic context (100kb) for each protein was obtained and analysed for presence of marker genes. We were looking for: 1) NRPS modules to see if the product of a potential nearby BGC is a peptide, 2) any TonB-dependent siderophore receptor, 3) FAAL domain, if acylation and subsequent deacylation are likely to occur during biosynthesis of a candidate; and 4) for C-domain specialised for accepting β-OH-Asp. The presence of a marker gene along with a putative peptide sequence was mapped onto the phylogenetic tree.

Siderophore-associated acylases formed a well-supported monophyletic lineage (cyanobacteriota 1, Fig. 5) with ∼47% of acylases associated with cyanochelin-like BGCs, clearly separated from cyanobacterial acylases that lack a siderophore BGC in their vicinity (cyanobacteriota 2). Interestingly, six acylases (∼8%) in the second lineage are found close to a NRPS BGC, however, subsequent inspection does not reveal other indications that the BGC would produce a siderophore.

Within the cyanobacteriota 1 lineage the overall branching pattern largely reflected the recently published genomic phylogeny of cyanobacteria (Strunecký et al. 2023), supporting the hypothesis of predominantly vertical inheritance of the acylase genes, likely as a part of the corresponding BGCs, with frequent secondary loss of the association to siderophore BGCs. Few notable exceptions to the general vertical evolution pattern are acylases from *Chroococcidiopsis thermalis* PCC 7203, *Gloeocapsa* sp. PCC 7428, *Gloeocapsopsis* dulcis AAB1 = 1H9 and *Synechocystis* sp. PCC 7509 that do not belong to Nostocales, however, all of them are representatives of the sister order-level lineage Chroococcidiopsidales (Strunecký et al. 2023). Possibly, this might be explained by an ancient horizontal gene transfer event between these two related lineages, or incomplete lineage sorting during the rapid ancient radiation event in the nitrogen fixing Nostocales/Chroococcidiopsidales lineage (Pardo-de la Hoz et al. 2023)..

The CcsQ resides in a branch of ∼13 acylases of which six are associated to BGCs, that contain all markers indicative of cyanochelin or cyanochelin-like siderophore product. Notably, only one acylase in the branch - WP_334311713.1 from *Kamptonema animale -* was found on a contig of sufficient length to safely state that none of the selected markers are in the vicinity. Additional two were found on contigs shorter than 10 kb and although none of the markers were detected. the sequence length does not allow to reliably exclude the possibility that the markers found in a relevant range. The remaining four acylases were found in a vicinity of BGCs that contained some of the selected features, yet again the BGC was incomplete due to contig boundary. The search strategy was set to reveal connection of acylases to putative siderophore BGCs and length filtering was not applied to exclude short nucleotide sequences. The presence of all selected markers even on a short stretch of DNA sequence provide enough evidence for the candidate acylase to be identified as a part of cyanochelin-like BGCs. However, for acylases, that were not found close to BGC or did not have all the markers we did not pursue extensive verification that they are not connected to siderophore BGC. Many of the acyalases that had some of the markers in the vicinity are actually likely to be a genuine part of a siderophore BGC that we could not confirm. The acylases of the CcsQ branch are often connected to peptides with conserved [Fatty_acid-X-Cys/Ser] motif on the N-terminus and possibly co-evolved with the first three NRPS modules.

**Fig. 4.**
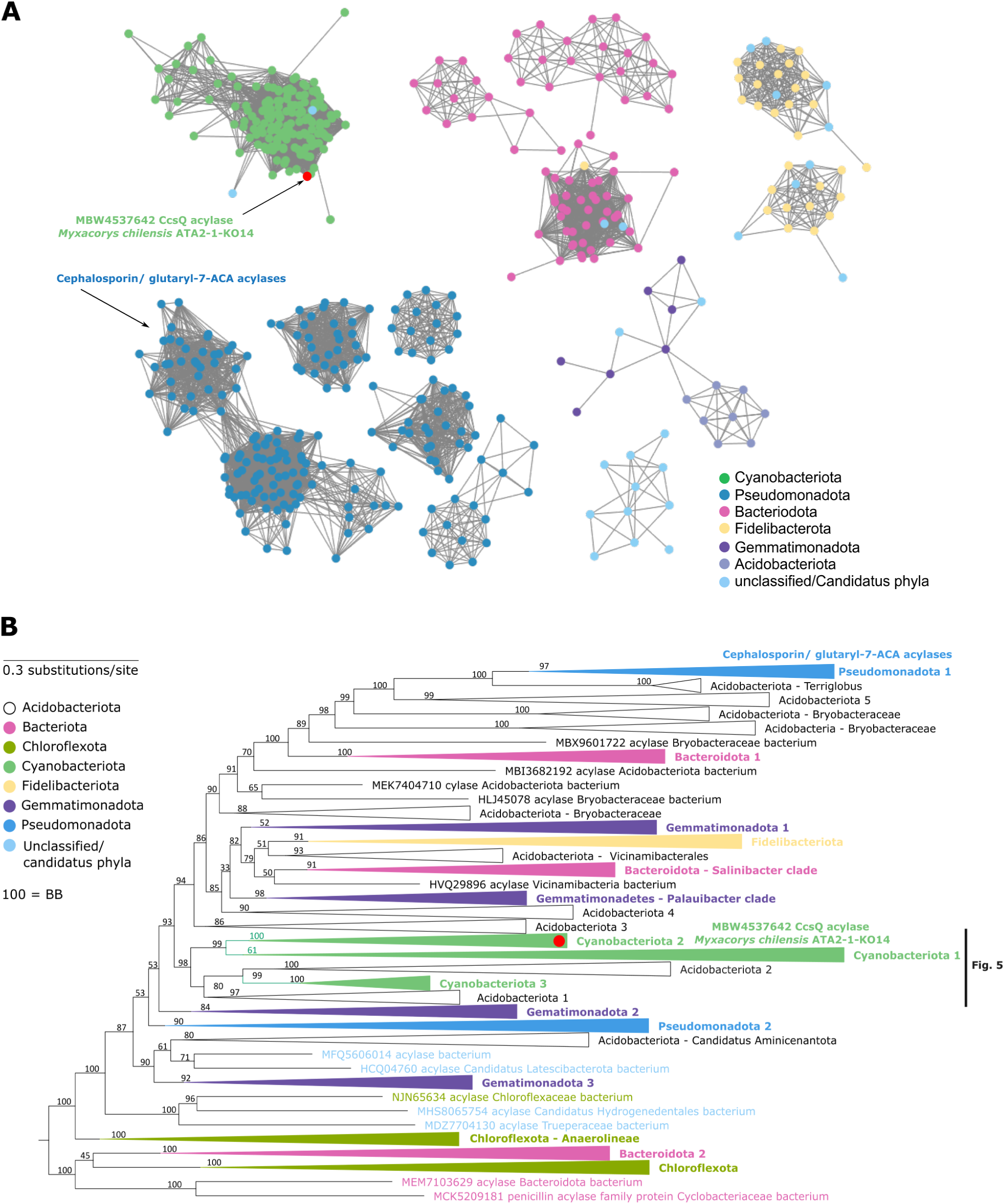
Phylogenetic analysis of CcsQ: **(A)** Sequence-similarity network of the best BLAST hits of CcsQ in bacteria. CcsQ is resolved within a coherent group of cyanobacterial acylase homologues. **(B)** Phylogenetic tree of CcsQ homologs

**Fig. 5.**
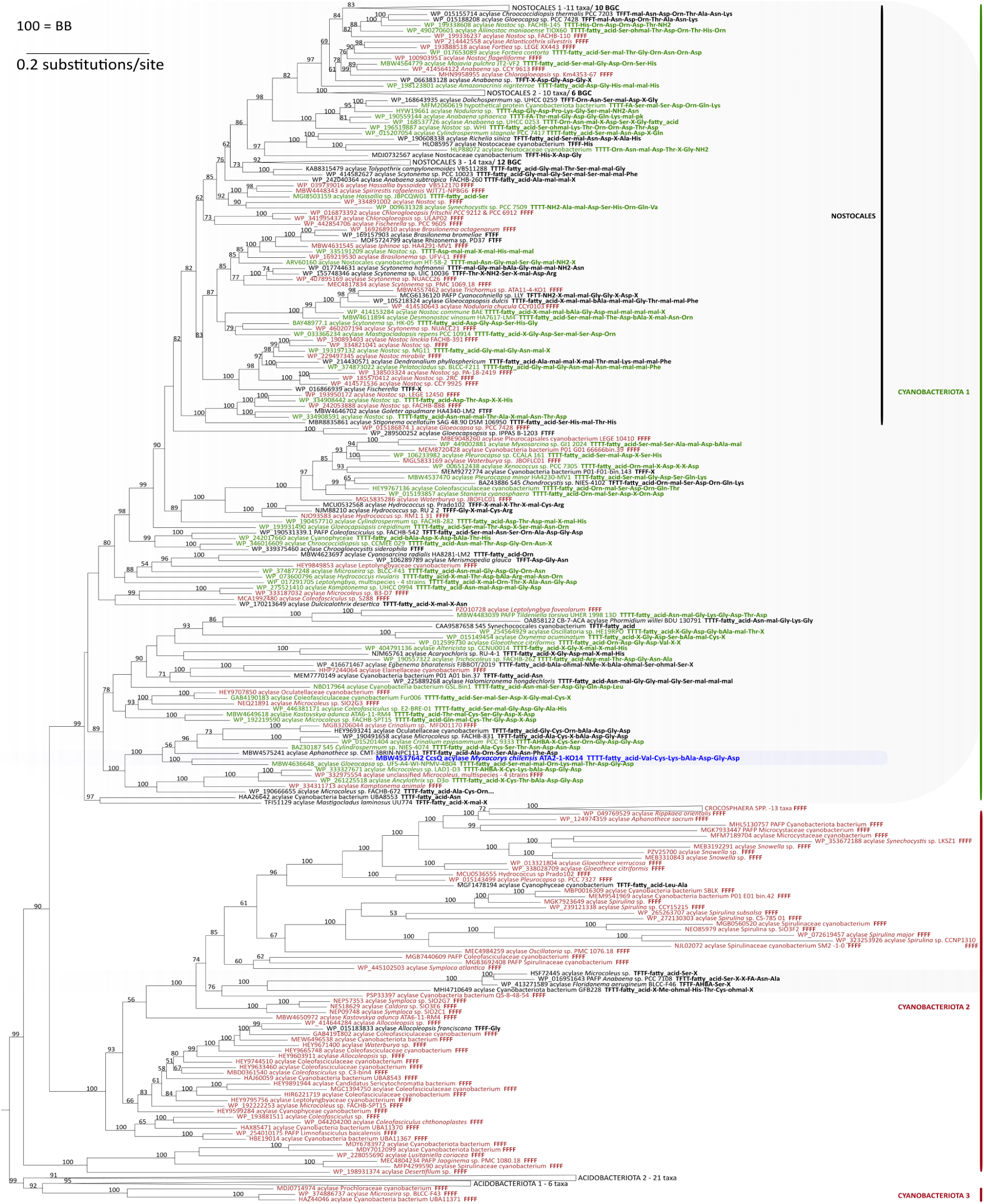
Reconstruction of the evolution of Cyanobacterial acylases; Siderophore-related cyanobacterial acylases, highlighted in grey form a well-defined group. CcsQ acylase is highlighted in blue. Major lineage with CcsQ shows majority of the included proteins to have association with BGCs and possibly siderophore biosynthesis (highlighted in green font).The other early branching lineages have vast majority of nodes not associated to putative siderophore BGCs (highlighted in red). Each node includes a protein id, protein description and strain of origin followed by a four letter code XXXX indicating the presence of 1) NPRS, 2) siderophore transporter, 3) FAAL domain, 4) β-OH-Asp-specific interface domain in the genomic vicinity of the acylase (T = true, F = false). If applicable, AntiSMASH-identified monomers of the final product are listed after the four letter code. Minor branches with multiple proteins from a taxonomically defined lineage are collapse and number of BGCs indicated. Abbreviations used in protein description: CB-7-ACA = 7-beta-4-carbaxybutanamido cephalosporanic acid, PAFP = penicillin acylase family protein, S45 = peptidase S45 penicillin amidase

## Discussion

The discovery of cyanochelin C, together with the corresponding BGC containing CcsQ, expands the still limited collection of cyanobacterial siderophores and sheds light on an interesting biosynthetic reaction that is likely not limited to this class of compounds. Genomic analyses have revealed numerous putative siderophore BGCs in cyanobacteria, however only few new cyanobacterial siderophores have been characterised recently, despite the availability of novel powerful techniques in natural product research and the well-known importance of iron for their cyanobacterial metabolism. Several factors may contribute to this limitation, including the slow growth of cyanobacteria, limited knowledge of their response to iron starvation, and complex interactions with other metals such as copper and zinc (Souza et al., 2025, Jeanjean et al 2008). Silent, putatively siderophore-encoding BGCs may therefore require an additional trigger to become active. Another drawback may come form presence of unknown tailoring enzymes the may considerably alter the properties of the compound. In this study, a prolonged iron starvation was sufficient to trigger compound production, however a tailoring enzyme, CcsQ, produced an initially unexpected pattern in HPLC/HRMS analyses. CcsQ shares functional similarities with known siderophore deacylases such as PvdQ, BntA, and MhtA/B; however, other, more unusual tailoring enzymes may still be undescribed in cyanobacterial BGCs. Such enzymes could substantially alter the chemical features of the final product, potentially leading to false negatives in screenings aimed at identifying silent BGCs.

Cyanochelin C shares multiple structural features with the other two cyanochelins described thus far (Fig. 6.; Galica et al 2021; Falcao et al 2025), including the N-terminal five-membered heterocycle and a motif involving two β-OH-Asp residues implicated in iron chelation. A key difference between cyanochelin C and cyanochelins A and B lies in the attachment of the fatty acyl (FA) moiety to the peptide backbone. In cyanochelins A and B, the FA moiety is first elongated and modified by a PKS module and then incorporated via a C–C bond. In contrast, cyanochelin C precursor carries a FA moiety directly linked to the N-terminal Val through a conventional amide bond. This distinction is reflected in the corresponding BGCs: whereas the FAAL domain in cyanochelins A and B is associated with PKS genes, in cyanochelin C it is fused directly to the NRPS assembly line. Notably, our previous bioinformatic analysis identified numerous cyanochelin-like BGCs with this FAAL-NRPS architecture, suggesting that cyanochelin C is the first characterised product of a biosynthetic strategy that may be widespread among cyanobacteria.

A further distinction concerns the metal-chelating moieties. While cyanochelin B contains two β-OH-Asp residues, cyanochelin A and C share an identical C-terminal peptide sequence, β-OH-Asp-Gly-DHABA. Efficient metal chelation requires correct positioning of the metal-binding residues. Both β-hydroxyaspartic acid and DHABA, each bearing two stereocenters, hence four possible configurations, need the proper absolute stereochemistry for binding. Resolving the configuration of β-hydroxyaspartic acid and DHABA found in cyanochelin C by Marfey’s method would require analytical standards of each configuration that are not easily accessible commercially. Instead, we relied on *J*-coupling analysis combined with bioinformatic analysis, an approach previously applied successfully to cyanochelin B. The stereochemical configuration of the C-terminal DHABA residue in cyanochelin A (C2) was not determined in the original study (Galica et al., 2021). Further examination of the NMR data, combined with bioinformatic analysis, suggests an *R* configuration at the β-carbon, matching the *R* configuration of the corresponding stereocenter of β-OH-Asp in cyanochelin B (C5; Falcão et al., 2025). In contrast, the corresponding stereocenter in cyanochelin C, the β-carbon (C2) of DHABA, is in the *S* configuration. The β-OH-Asp of cyanochelin C is in the L-*erythro* configuration (*8S,9R*), which, according to a comprehensive study by Reitz et al. (2019), is less common and has so far been identified only in imaqobactin (Reitz et al., 2019; Robertson et al., 2018). Genome analysis suggests that several other siderophores, such as cupriachelin, histicorrugatin, and delftibactin, may also contain L-*erythro* β-OH-Asp, but their stereochemistry has not yet been experimentally resolved. Presumably, all chiral configurations of β-OH-Asp can be employed depending on the surrounding structural context, with the conformation finely tuned to achieve proper positioning for metal chelation by other structural elements such as D-Lys or β-Ala. The N-terminal heterocycle is present in all cyanochelins described thus far: an oxazole in cyanochelin A and C, and a thiazole in cyanochelin B. Whether this heterocycle contributes directly to iron chelation remains an open question. In the bacterial siderophore mycobactin, an oxazole heterocycle participates in chelation together with a phenolate moiety (Snow 1970). In cyanochelin A and B, hydroxyl substituents in the vicinity of the heterocycle might play an analogous role. Curiously, cyanochelin C lacks any hydroxyl substitution near its oxazole heterocycle, raising the question of whether the heterocycle contributes to chelation in this compound at all.

**Fig. 6.**
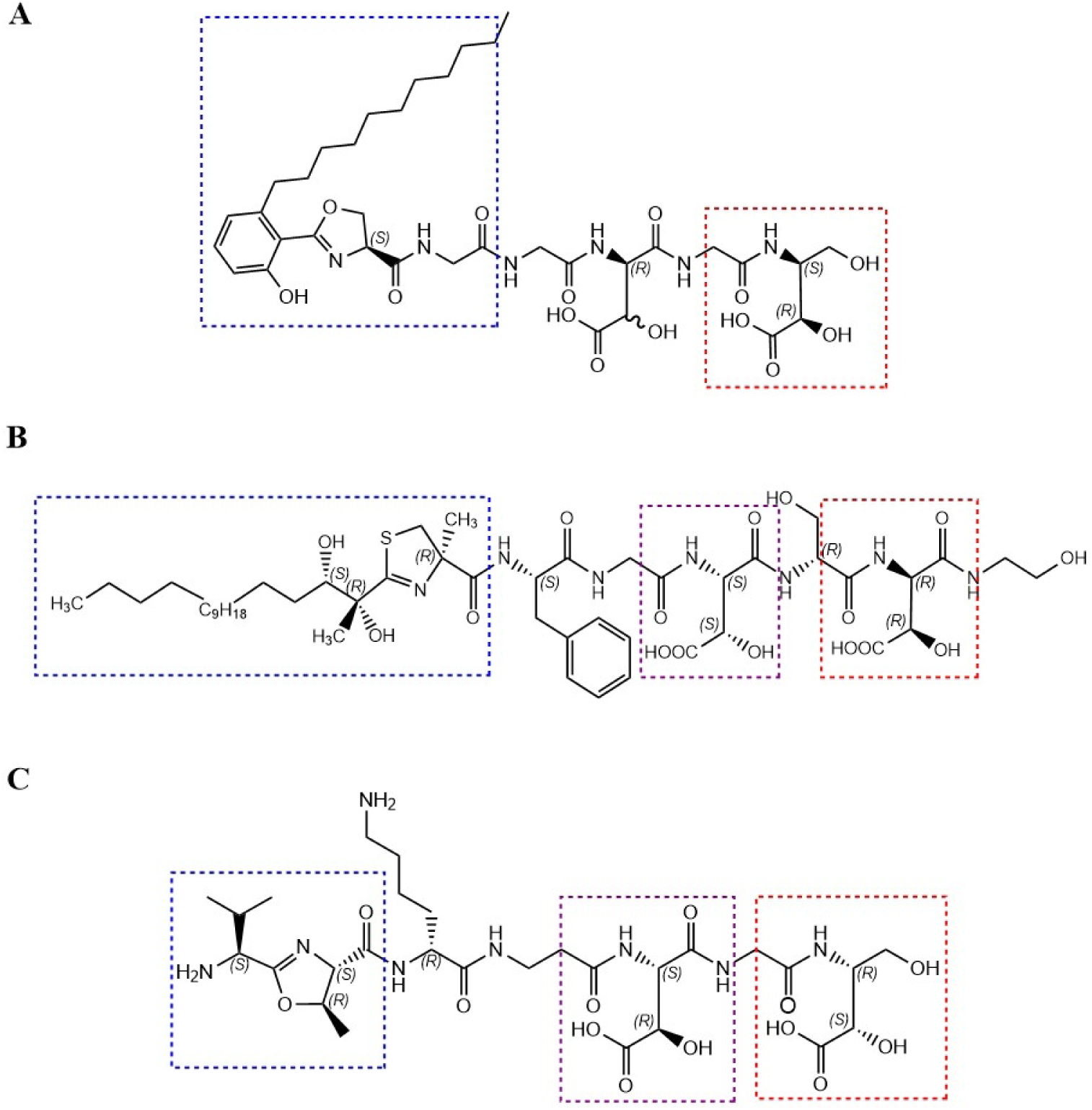
Comparison of structural motifs across cyanochelins A, B and C.

Acylation is a frequent modification of cyanobacterial natural products, and FAAL domain is a common starter module in cyanobacterial NRPS BGCs (Fewer et al., 2021; Galica et al., 2017; Mareš et al., 2019). This also holds true for siderophores in general, and both cyanochelin A and B are acylated. As mentioned earlier, siderophore variants often differ in the presence and/or length of an acyl residue. Some, such as pyoverdines, mature via acyl cleavage by a dedicated acylase (PvdQ); others, such as marinobactins and aquachelins, occur as suites of variants differing in acyl-chain length, with de-acylated forms arising optionally depending on the presence of a compatible acylase in the producer or a cohabiting strain (Kem et al., 2015; Gauglitz et al., 2014). For products in which hydrophobicity is not essential, initial acylation may improve docking to biosynthetic enzymes and increase the efficiency of subsequent biosynthetic steps. For others, deacylation may fine-tune diffusibility or mediate a switch between functions in the cell’s immediate vicinity versus its wider surroundings. For cyanochelin C, the presence of *ccsQ* in the BGC, possibly sharing the promoter region with the NRPS biosynthetic genes, suggests that the deacylation of the preproduct is by design. Interestingly, in contrast to cyanochelin A and B, where the peptide bond connecting the acyl residue to the peptide core is converted and becomes a part of the heterocycle, in cyanochelin C the methyloxazoline is at position 2, and the acyl chain is connected to N-terminal valine by a regular peptide bond (Fig. 6). This makes the enzymatic deacylation of cyanochelin C possible, while the chemical structure of the other two described cyanochelins is unlikely to be cleaved by a CcsQ or its homolog. Either the proper function of cyanochelin A and B depends on the acylation, or the switch to hydrophilic siderophore is facilitated by photolysis-mediated removal of acyl chains as observed in cyanochelin B. It is worth noting that while previously inspected cyanochelin A and B were predominantly associated with biomass, cyanochelin C is almost exclusively found in spent media at high titres.

The closest experimentally characterised enzymes to CcsQ are cephalosporin/glutaryl-7-cephalosporanic acid (GL-7-ACA) acylases, which industrially convert over 1,000 tons of cephalosporin C to 7-ACA annually, a key step in low-waste antibiotic production (Lin and Kuck, 2022; Gröger et al., 2017). These enzymes are structurally well characterized, including several crystal structures with mutational and kinetic data (Kim et al., 2000; Kim et al., 2003; Fritz-Wolf et al., 2002; Pollegioni et al., 2005; PDB 1GK0, 1GK1), which should facilitate modelling of CcsQ. They hydrolyse amide bonds linking small substituents to β-lactam nuclei (Oinonen and Rouvinen, 2000; Velasco-Bucheli et al., 2020), and their compact, polar substrate-binding pockets favour small, carboxylate-rich substrates over long hydrophobic acyl chains (Kim et al., 2000; Velasco-Bucheli et al., 2020). Like PvdQ, BntA, and MhtA, they belong to the Ntn-hydrolase superfamily. Phylogenetically, CcsQ clusters with GL-7-ACA/cephalosporin acylases rather than with PvdQ-like AHL acylases, despite its PvdQ-like presumed function, possibly reflecting selection for interaction with a polar peptide core rather than for a hydrophobic acyl-binding tunnel.

Because research on GL-7-ACA acylases has been largely application-driven, the natural roles of their relatives remain understudied. Related AHL acylases participate in siderophore maturation and quorum sensing/quenching; for example, AiiC from *Anabaena* sp. PCC 7120 interferes with quorum sensing (Romero et al., 2008) and, although not BGC-adjacent, could still influence siderophore biosynthesis. *Anabaena* produces schizokinen and a putative second uncharacterised siderophore (Jeanjean et al., 2008). The schizokinen BGC contians an acyltransferase that could produce synechobactins by acylation of schizokinen. Hence schizokinen may arise directly, as an unacylated product of the BGC, or via synechobactin deacylation by AiiC, as seen for marinobactins.

Phylogenetic analysis of cyanobacterial acylases suggests deep branching into two main lineages, one of which includes CcsQ and is clearly associated with secondary metabolites, most frequently with putative siderophores. CcsQ like acylases are present across major cyanobacterial lineages in a manner that resembles distribution of other secondary metabolite tailoring enzymes. A dedicated bioinformatic screening would be required to determine the frequency of acylase occurrences in siderophore BGCs, however, inspection of our previous data found an acylase in 22 out of 31 cyanochelin-like BGCs (Galica et al 2021). Siderophore deacylation is likely common in cyanobacteria, though further examples and substrate-range data are needed to confirm a dedicated maturase role.

Ecologically, siderophore diffusibility shapes iron-binding efficiency, competition, piracy resistance, and environmental adaptation. Having previously examined this for cyanochelin B (Falcão et al., 2025; Falcão et al., manuscript in preparation), cyanochelin C now allows us to probe the effect of acylation on these interactions. Since CcsQ homologs also cleave AHLs, iron-starved producers may simultaneously deploy cyanochelin C and disrupt bacterial quorum sensing, amplifying their ecological impact.

Cyanochelin C thus expands the cyanobacterial siderophore catalogue with a compound enabling comparisons across acylation states and chelating-residue stereochemistry, while characterization of CcsQ reveals a likely widespread, previously overlooked biosynthetic step with implications for microbial communication and biotechnology, illustrating how siderophore research can uncover new enzyme families relevant to cyanobacterial ecology and biotechnology more broadly.

## Methods

### Strains and cultivation

*Myxacorys chilensis* was kindly provided by Jeffrey R. Johansen. The strain was first primed in BG-11 medium and further iron starved in BG-11 medium without the addition of ferric Ammonium Citrate at small scale (20 ml). After complete starvation and siderophore production, the starved biomass was upscaled to 2 litres and incubated at ambient temperature (25 ± 1°C) with continuous dispersed light and constant bubbling. The culture was checked regularly for siderophore production using the CAS assay and the HPLC-HRMS method mentioned below.

### Isolation of Cyanochelin C

Iron-deprived and CAS-positive culture of *Myxacorys chilensis* was centrifuged at 2000 g for 10 minutes, room temperature to remove the biomass. Subsequently the supernatant was subjected to solid phase extraction (SPE) on C-18 reverse phase Supelco Discovery tube (10g, 60mL). The column was activated by methanol and briefly flushed by distilled water to stabilise before application of the spent media. After passing of the culture media the column was washed with distilled water to get rid salts and any polar impurities. The compounds retained were eluted with 35mL methanol. The extract was then subjected to rotary evaporation and reconstituted in water with 10% acetonitrile (900 µL). The redissolved extract was subjected to preparative reverse phase HPLC on an Agilent Eclipse XDB-C8 column (9.4 x 250 mm, 5 µm). Water (A) and ACN (B) were used as mobile phase in the following gradient: 0-4 min 10% B, 4 min 10% B, 25 min 100% B, 28 min 100% B, 29-30 min 10% B at flow rate 5 ml/min and max pressure 380 bar. The multiwavelength detector (MWD) was set to 254 nm and used for detection of Cyanochelin C. Cyanochelin C was eluted at 12 min. The content of the collected fractions was analysed by HPLC-HRMS/MS as described below.

### HPLC-HRMS/MS

Content of Cyanochelin C cultures and fractions was analyzed using the Dionex Ultimate 3000 HPLC system coupled to Bruker Impact HD II mass spectrometer equipped with electrospray ionization. Separation was performed using Waters MaxPeak XBridge Premier BEH C18 chromatographic column (130 Å, 2.5 µm; 50 × 2.1 mm). Acetonitrile (solvent A) (LCMS grade, Honeywell) and water (solvent B; LCMS grade, Honeywell), supplemented with trifluoroacetic acid (≥99.0%, CHROMASOLV for LCMS, Fluka, Avantor, cat. no. 302031) at concentrations of 0.002% and 0.1% vol/vol, were used as mobile phases. The column was eluted according to the method described by Falcao et al, 2025. Spectra were collected in the range of 50 to 1,490 m/z with a spectral rate of 2 Hz.

### Structural characterization of cyanochelin C

1D NMR and 2D NMR experiments were carried out at 25°C on a Bruker AvanceNeo 700 MHz spectrometer (Billerica, MA, US) equipped with a triple resonance CHN cryoprobe. The ^1^H NMR spectrum was acquired in D_2_O/H_2_O (9:1) (Sigma Aldrich, Milan, Italy) containing 20 mM phosphate buffer, whereas all 2D experiments were acquired in D2O (Sigma Aldrich, Milan, Italy); chemical shifts were referenced to the residual solvent signal (D_2_O: δ_H_ 4.79 ppm). The HSQC spectra were optimized for ^1^*J*_C,H_ = 145 Hz. The HMBC experiments for ^2,3^*J*_C,H_ = 8 Hz. Abbreviations for signal couplings are as follows: s = singlet, d = doublet, br.d = broad doublet, dd = doublet of doublets, ddd = doublet of doublets of doublets, t = triplet, q = quartet, quint= quintet, m = multiplet.

### Advanced Marfey’s analysis

The cyanochelin C (10 μg) was hydrolyzed with 600 μL of 6N HCl/AcOH (1:1) at 120 °C for 18 h. The residual HCl fumes were removed under an N_2_ stream. The hydrolysate was then dissolved in TEA/acetone (2:3, 200 μL) and the solution was treated with 200 μL of 1% 1-fluoro-2,4-dinitrophenyl-5-L-alaninamide (L-FDAA) in CH_3_CN/acetone (1:2) (Esposito et al, 2016). The mixture was dried, and the resulting L-DAA derivatives of the free amino acids were redissolved in MeOH (200 μL) for subsequent analysis. Authentic standards of L-Lys, L-Val and L-Thr were treated with L-FDAA and L-FDAA as described above and yielded the L-DAA and D-DAA standards. Marfey’s derivatives of cyanochelin C were analyzed by using a Thermo Scientific Orbitrap Exploris 120 high-resolution ESI mass spectrometer (Thermofisher, Waltham MA, USA), integrated with a Thermo U3000 HPLC system (Thermofisher, Waltham MA, USA), and their retention times were compared with those from the authentic standard derivatives. A 5 μm Kinetex C18 column (50 × 2.10 mm) maintained at 25 °C was eluted at 200 μL min^−1^ with 0.1% HCOOH in H_2_O and ACN. The gradient program was as follows: 5% ACN 3 min, 5–75% ACN over 30 min, 75-95% ACN over 1 min and 95% ACN 6 min.

Mass spectra were acquired in positive ion detection mode. MS parameters utilized a spray voltage of 4.8 kV, a capillary temperature of 285 °C, a sheath gas rate of 32 units N _2_ (ca. 230 mL/min), and an auxiliary gas rate of 15 units N_2_ (ca. 150 mL/min). The MS method involved four HRMS/MS scans after each full MS scan for the four most intense ions detected in the spectrum (data-dependent acquisition mode, DDA). The *m/z* range for data-dependent acquisition was set between 135 and 2000 amu with resolution set to 120,000. HRMS/MS scans were obtained for selected ions with CID fragmentation using an isolation width of 1.5, a normalized collision energy of 30, an activation Q of 0.250, and an activation time of 30 ms. Data analysis was conducted using Thermo Xcalibur software (version 2.2 SP1 build 48).

### Phylogenetic analyses and Sequence-Similarity Network

A Sequence-Similarity Netwok (SSN) was created using the online EFI – Enzyme Similarity Tool with UniProt version 2025-03 (Gerlt et al. 2015). Initial BLAST analysis employed CcsQ (NCBI: MBW4537642.1) as a query using the UniProt database and E-value of 20 for BLAST hit retrieval. After visual inspection of the percent identity vs. BLAST alignment score plot, an alignment score of 250 (corresponding to ∼50% percent identity) was used as a threshold for connecting proteins in the network. The SSN was visualized using Cytoscape v. 3.10.3 (Shannon ert al. 2003), nodes were classified based on Phylum name; singletons and small groups of proteins (<10 nodes) were omitted from the final graph for clarity.

To determine the evolutionary relationship of cyanochelin C acylase CcsQ (MBW4537642) to known enzymes, a set of representative amino-acid sequences of experimentally characterized N-terminal nucleophile hydrolases was assembled based on literature search and custom BLASTp analysis of CcsQ and pyoverdine acylase PvdQ (AAG05773). The resulting set of proteins was broadened by several more distant homologues according to Velasco-Bucheli et al. (2020). Protein sequences were aligned using MAFFT v. 7.490 (Katoh and Standley 2013) with default settings, and gap regions were removed manually. The Maximum Likelihood (ML) phylogenetic reconstruction was performed using RaxML v. 8 (Stamatakis 2014) with the GAMMA LG model and 1,000 rapid bootstrap repetitions mapped on the best-scoring ML tree.

### Protein modelling

Sequence of CcsQ was submitted to AlphaFold server (https://alphafoldserver.com/) which runs AlphaFold3. A peptide sequence Leu-Ser-Lys-Ala-Asp-Ala-Asp mimicking the assumed siderophore without modification of amino acids was submitted as an additional protein chain and the myritstic acid was added as a ligand. The obtained results were downloaded and visualised using PyMOL Molecular Graphics System Version 3.1.3 Copyright (C) Schrödinger, LLC. (https://pymol.org/), with highlighting the amino acids in the active site as found in Drake and Gulick 2011.

### Phylogenetic analysis of CcsQ-like N-terminal nucleophile hydrolases in cyanobacteria

Amino acid sequence of CcsQ from *M. chilensis* ATA2-1-KO-14 (MBW4537642.1,) was used as query sequence against clustered protein database in a BLASTp analysis via online NCBI BLAST suite. The top 1 000 BLASTP hits were retrieved and manually curated by removing partial, unusually long, and highly divergent sequences. Multiple sequence alignment was generated using MAFFT (Katoh and Standley 2013) implemented through the Galaxy platform (Jalili et al. 2020). The alignment was subsequently inspected and manually curated using the MUSCLE (Edgar 2004) algorithm implemented in MEGA (Tamura et al. 2021). Poorly aligned terminal regions and two ambiguously aligned internal regions were excluded from further analyses. The final alignment comprised 642 amino acid positions from 928 protein sequences. No outgroup taxa were chosen, since we are not sure about the evolution direction. Phylogenetic relationships were inferred using the maximum likelihood (ML) method implemented in IQ-TREE (Nguyen et al. 2015). The best-fitting amino acid substitution model was selected using ModelFinder (Kalyaanamoorthy et al. 2017) also implemented in IQ-TREE. Branch support was assessed using the ultrafast bootstrap (BB) approximation with 2 000 replicates.

For detailed phylogenetic analysis, only cyanobacterial sequences (259 taxa) and their closest Acidobacteriota homologues (31 sequences), selected as the outgroup based on the previous comprehensive acylase phylogeny, were included. Multiple sequence alignment and curation were performed as described above. The final dataset comprised 292 taxa and 642 aligned amino acid positions. The dataset was analysed by ML method as described above.

For each of 259 cyanobacterial proteins, identified as CcsQ homologues, a nucleotide sequence encoding the protein and surrounding genomic region (50kb upstream and downstream) were obtained from NCBI. The sequence was analysed using AntiSMASH to detect NRPS siderophore-encoding BGCs and the substrate specificity of adenylation domains found in the cluster. To further evaluate if the acylase may be associated with a cyanochelin-like siderophore BGC the genomic context of each acylase was furhter inspected for the presence of 3 markers: TonB-dependent siderophore receptors (TBDT), fatty acid-AMP ligase domain (FAAL) and β-OH-Asp-specific condensation domain (DOXC). BLAST database was constructed from all proteins encoded within the obtained nucleotide sequences and searched for proteins similar to PSSMs of TBDT, FAAL and DOXC, scoring over a manually determined threshold. Presence of NRPS, TBDT, FAAL, DOXC and NRPS-recognized monomers was mapped manually to individual acylases in the phylogenetic tree.

## Supporting information

Table S1: Analysis of transporter cassette

Figures S1-S10 - NMR spectra of Cyanochelin C

Fig S11 - full phylogenetic tree of cyanobacterial acylases

## Data availability

The NMR data used for structure elucidation of cyanochelin C have been deposited in the Natural Products Magnetic Resonance Database (NP-MRD; www.np-mrd.org) and can be found under accession number NP0354349.

## Acknowledgements

This work was supported by Czech Science Foundation (22-05478S - Iron monopolization versus community service: the two faces of cyanobacterial beta-hydroxy aspartate lipopeptides). Additional support was provided by the Grant Agency of the University of South Bohemia (GAJU), project no. 126/2024/P, “Do heterotrophic bacteria hijack cyanobacterial siderophores via specific transport mechanisms?” (PI: Berness Peter Falcao). Additional support was provided by OP JAK project “Photomachines” Reg. No CZ.02.01.01/00/22_008/0004624 (Czech Ministry of Education, Youth, and Sports (MEYS)), financed by the Ministry of Education, Youth and Sports. The authors acknowledge the support of NBFC to the University of Napoli Federico II, Department of Pharmacy, funded by the Italian Ministry of University and Research, PNRR, Missione 4 Component 2, “Dalla ricerca all impress”, Investimento 1.4, Project CN00000033. The research was also funded as part of the program “Finanziamento della Ricerca di Ateneo” (FRA) 2022, by Università degli Studi di Napoli Federico II, with the contribution of Compagnia di San Paolo. Lenka Marešová Štenclová was supported by Rising-Star-Fellowship of the Department of Biology, Chemistry, Pharmacy at Freie Universität Berlin.

