## Supplementary material for "Structural and Stereochemical Elucidation of Cyanochelin C, a Siderophore Associated with Novel Class of Cyanobacterial Acyl Hydrolases": Table S1: Analysis of transporter cassette

| **Position** | ***Gene Name*** | **size (AA)** | **BLAST Annotation** | ***M. chilensis*** **(JAHHHP010000001) Protein ID:** | **M. californica (JAFJZQ010000037) Protein ID:** | **Functional homology** | ***Leptolyngbya*** **sp. NIES 3755 (AP017310) Protein ID:** | **Coverage/ Identity [%]** |
|---|---|---|---|---|---|---|---|---|
| **-8** | ***CctC*** | 436 | MFS transporter | MBW4537650.1 | MBW4417605.1 | ***cctC*** | BAU16019.1 | 99/47 |
| **-7** | ***cctB1/J*** | 355 | iron-siderophore ABC transporter substrate-binding protein | MBW4537649.1 | MBW4417606.1 | **paralogue *cctB1*** | BAU16018.1 | 82/31 |
| **-6** | ***cctB2/B*** | 347 | iron-siderophore ABC transporter substrate-binding protein | MBW4537648.1 | MBW4417607.1 | ***cctB2*** | BAU16018.1 | 94/41 |
| **-5** | ***cctD*** | 325 | sucrase ferredoxin | MBW4537647.1 | MBW4417608.1 | ***cctD*** | BAU16020.1 | 72/30 |
| **-4** | ***cctA*** | 843 | TonB-dependent siderophore receptor | MBW4537646.1 | MBW4417609.1 | ***cctA*** | BAU16017.1 | 77/26 |
| **-3** | ***cctE*** | 354 | iron ABC transporter permease | MBW4537645.1 | MBW4417610.1 | ***cctE*** | BAU16021.1 | 99/57 |
| **-2** | ***cctF*** | 346 | iron ABC transporter permease | MBW4537644.1 | MBW4417611.1 | ***cctF*** | BAU16022.1 | 99/58 |
| **-1** | ***cctI*** | 277 | ABC transporter ATP-binding protein | MBW4537643.1 | MBW4417612.1 | ***cctI*** | BAU16025.1 | 88/32 |

**Table S1:** Analysis of transporter cassette found in the vicinity of Cyanochlin C BGC
