## Supplementary material for "Structural and Stereochemical Elucidation of Cyanochelin C, a Siderophore Associated with Novel Class of Cyanobacterial Acyl Hydrolases": Figures S1-S10 - NMR spectra of Cyanochelin C

**Supporting Material Draft**

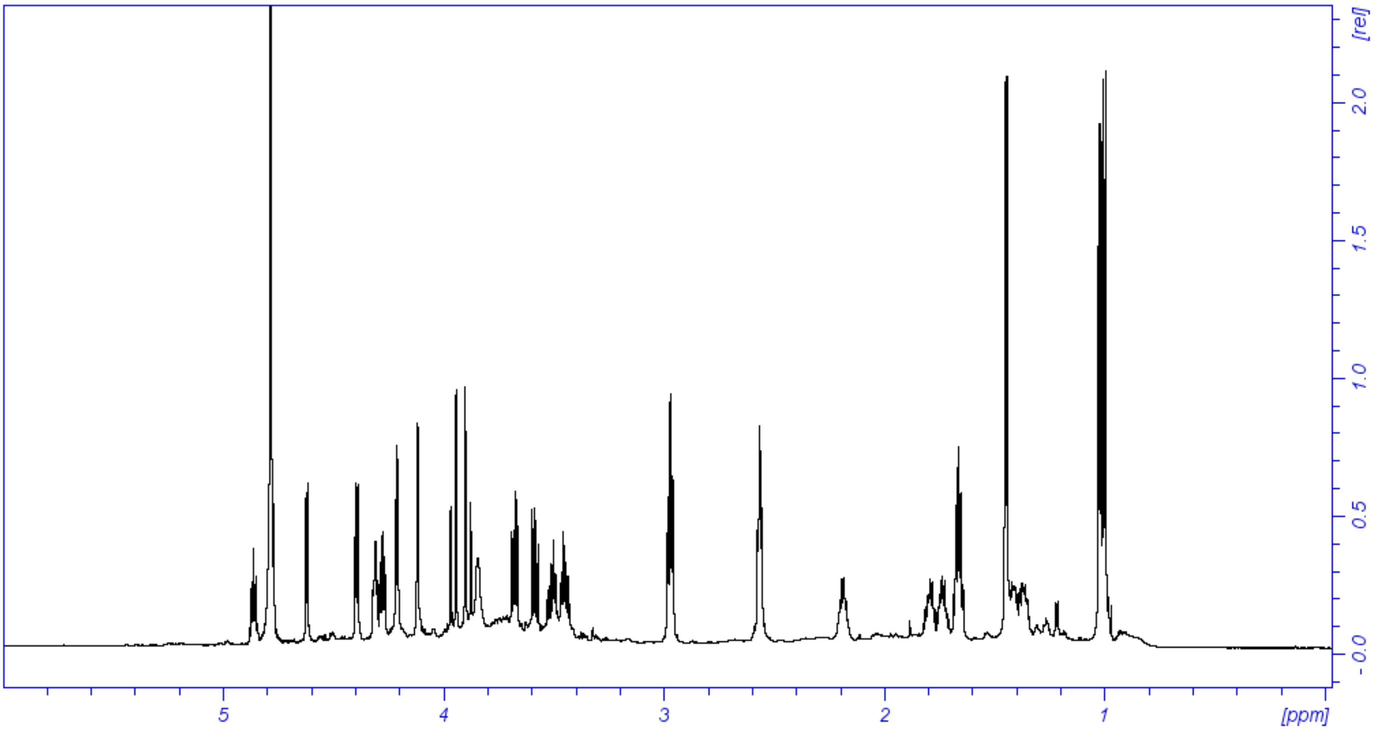


**Figure S1:** $^{1}$H-NMR spectrum of cyanochelin C (700 MHz, $D_2O$, 298K)

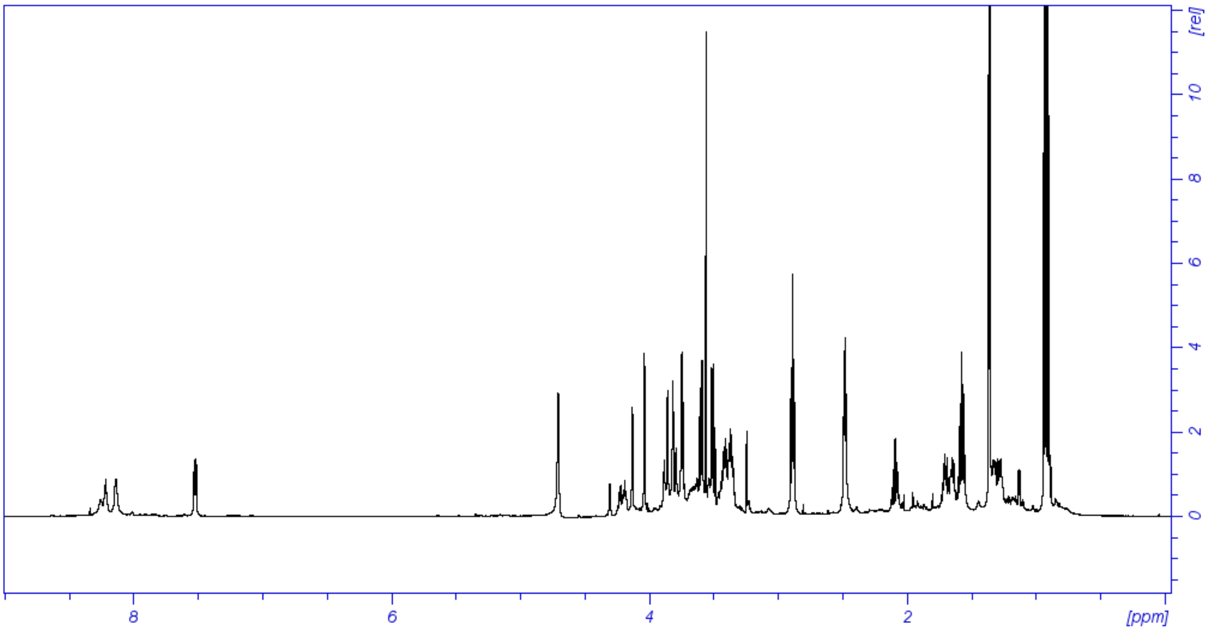


**Figure S2:** $^{1}$H-NMR spectrum of cyanochelin C (700 MHz, $D_2O/H_2O$ 9:1, 398K)

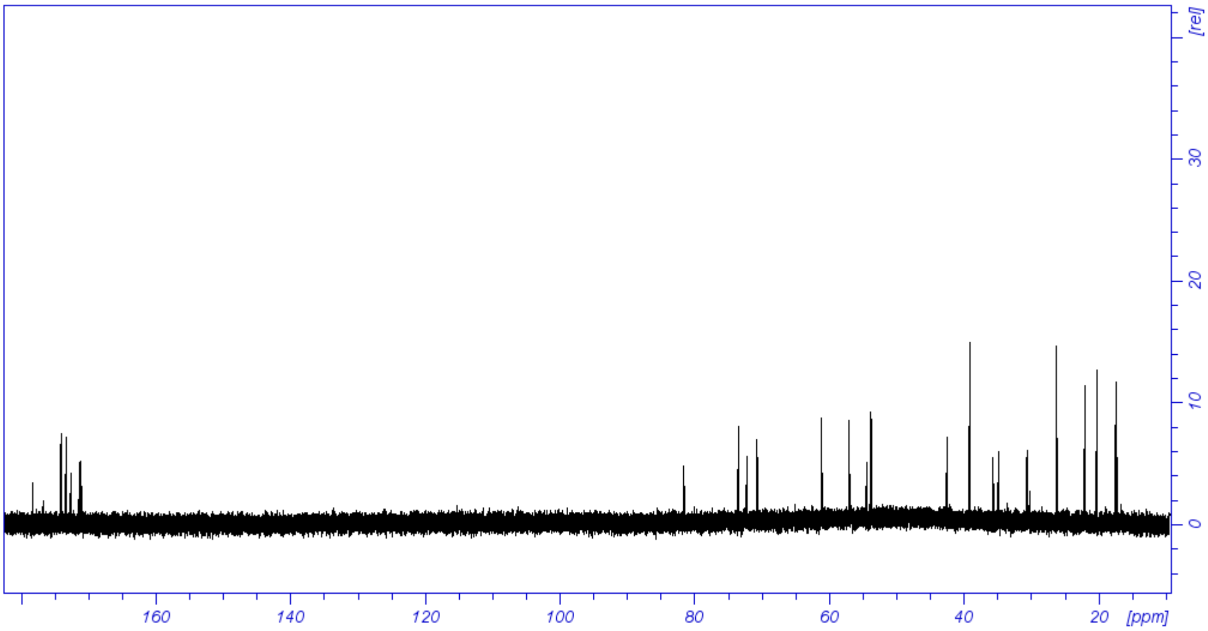


**Figure S3:** $^{13}$C-NMR spectrum of cyanochelin C (700 MHz, $D_2O$, 298K)

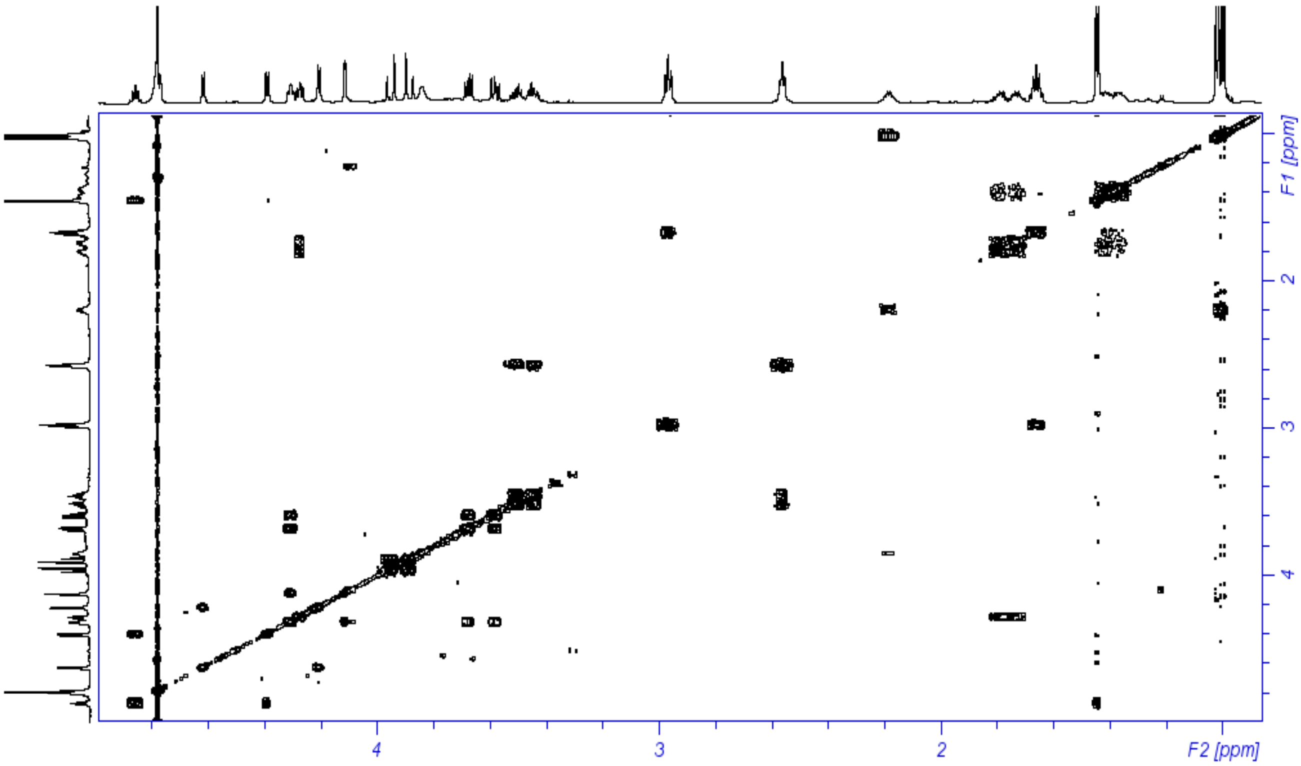


**Figure S4:** COSY spectrum of cyanochelin C (700 MHz, $D_2O$, 298K)

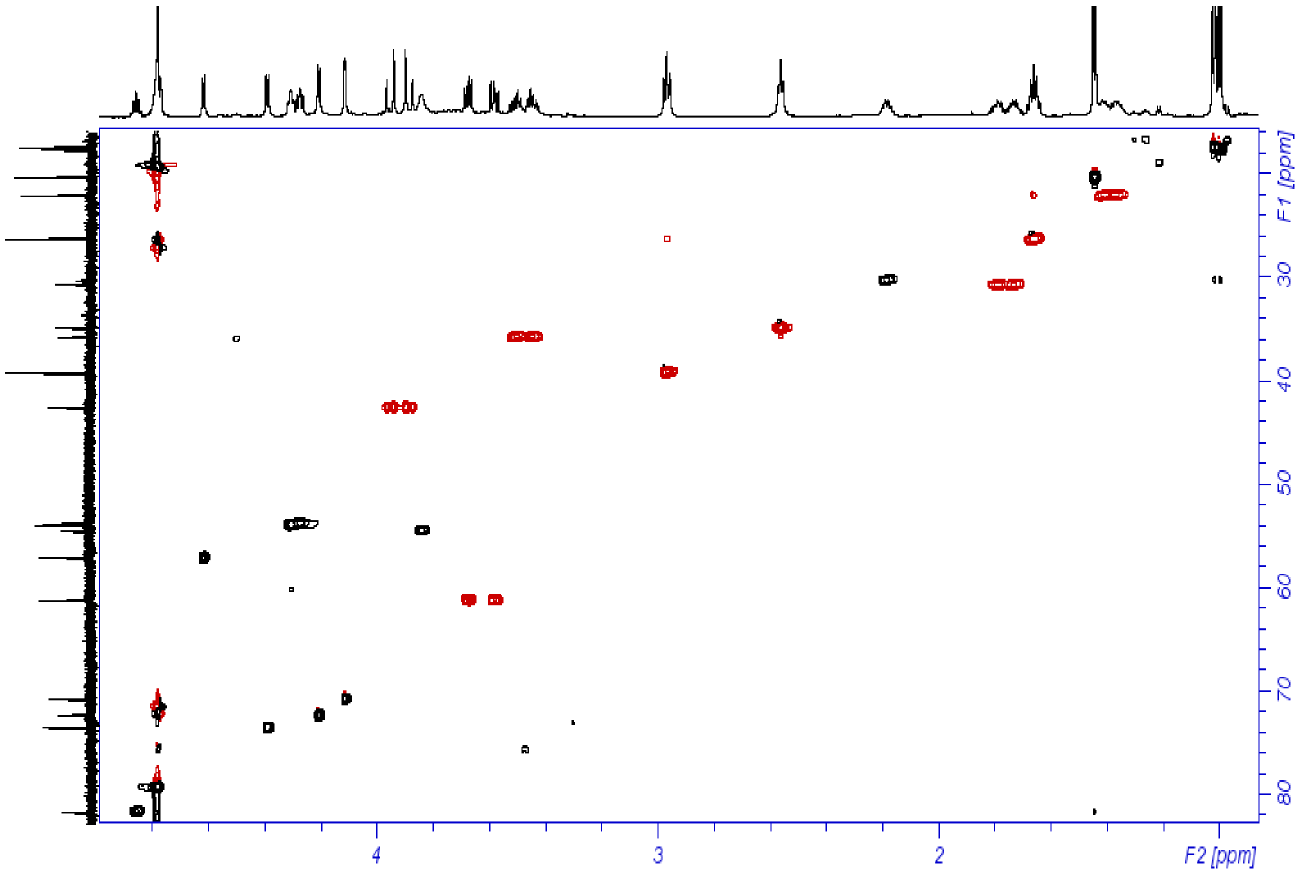


**Figure S5:** HSQC spectrum of cyanochelin C (700 MHz, $D_2O$, 298K)

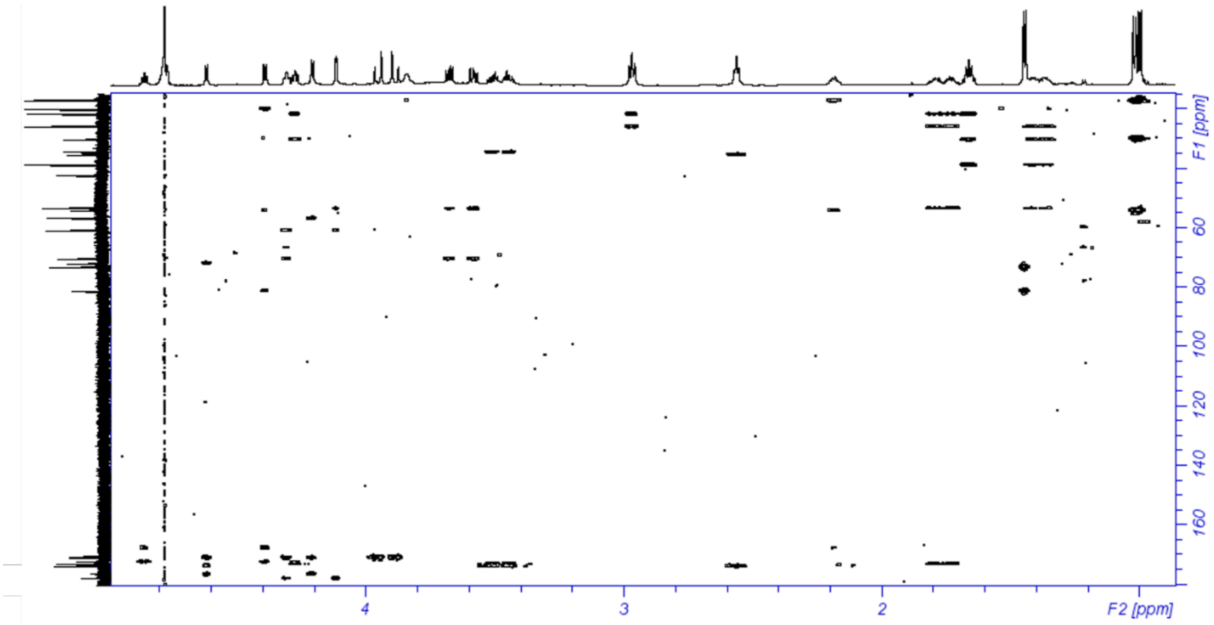


**Figure S6:** HMBC spectrum of cyanochelin C (700 MHz, $D_2O$, 298K)

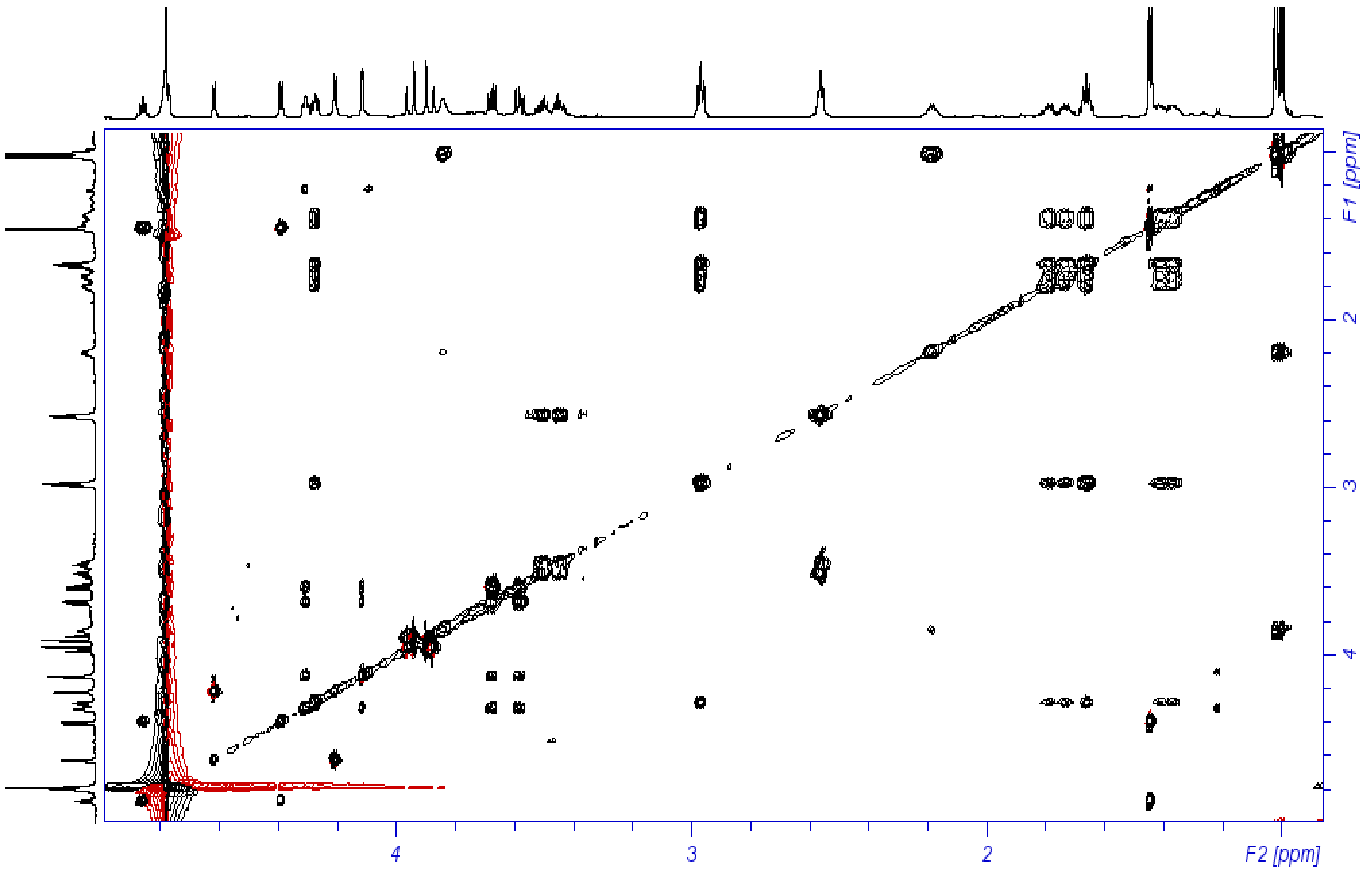


**Figure S7:** TOCSY spectrum of cyanochelin C (700 MHz, $D_2O$, 298K)

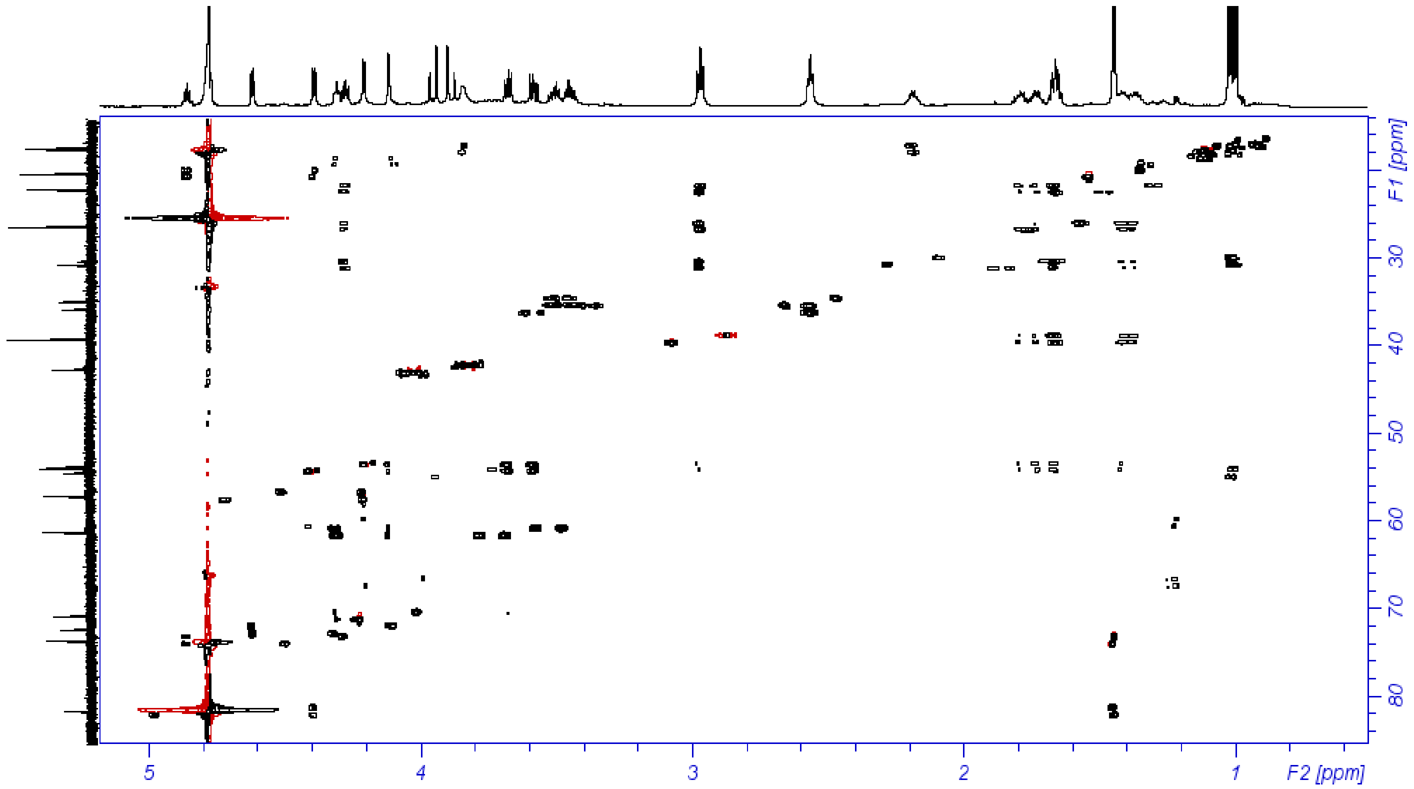


**Figure S8:** HSQC-HECADES spectrum of cyanochelin C (700 MHz, $D_2O$, 298K)

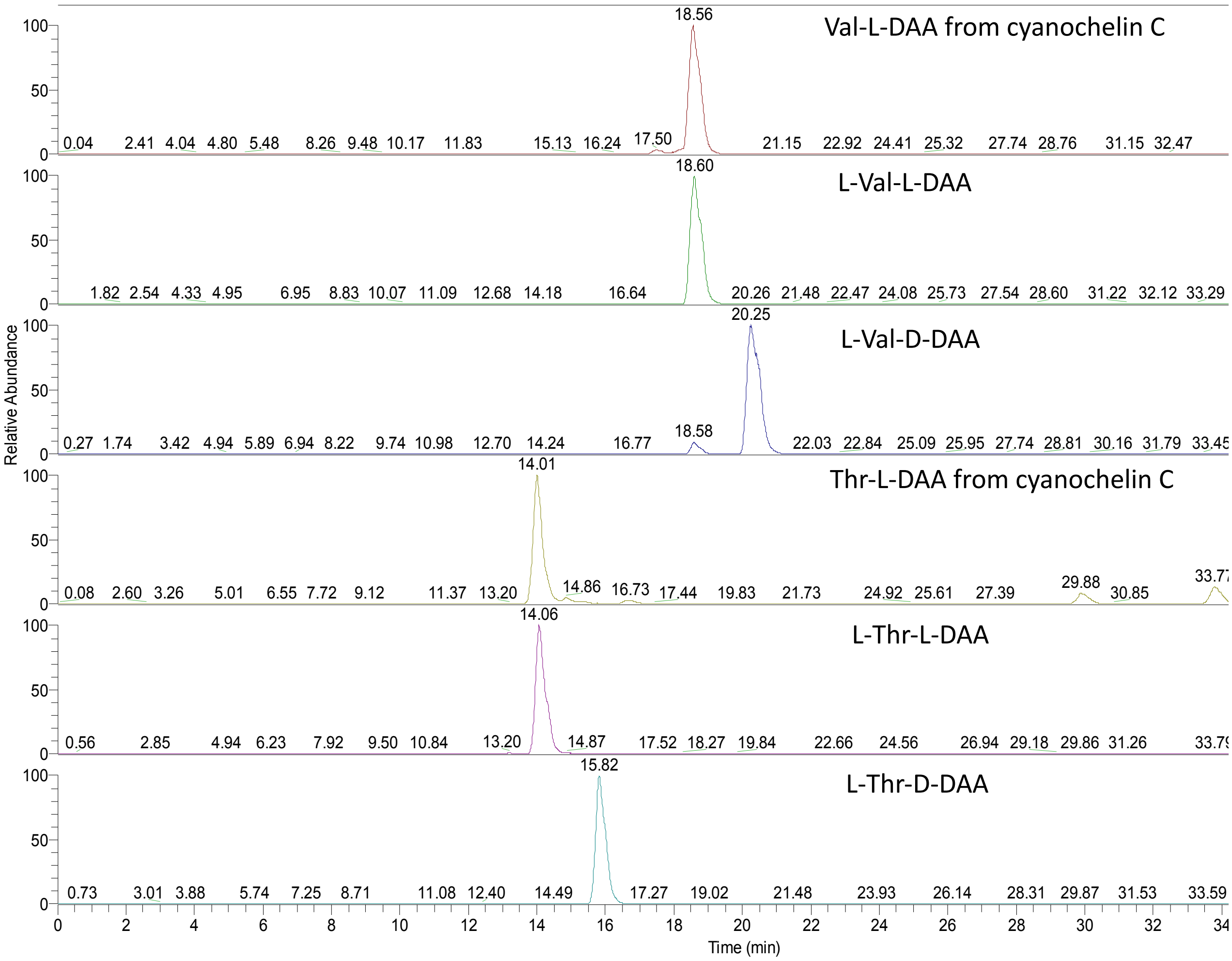


**Figure S9:** LC-MS-enhanced Marfey's analysis of cyanochelin C. Extracted-ion chromatograms at *m/z* 370.1357 of the L-FDAA derivative from the hydrolysis of cyanochelin C and of D-and L-FDAA derivatives of L-Val. Extracted-ion chromatograms at *m/z* 372.1150 of the L-FDAA derivative from the hydrolysis of cyanochelin C and of D-and L-FDAA derivatives of L-Thr.

**Figure S10:** LC-MS-enhanced Marfey's analysis of cyanochelin C. Extracted-ion chromatograms at *m/z* 399.1623 of the L-FDAA derivative from the hydrolysis of cyanochelin C and of D-and L-FDAA derivatives of L-Lys.

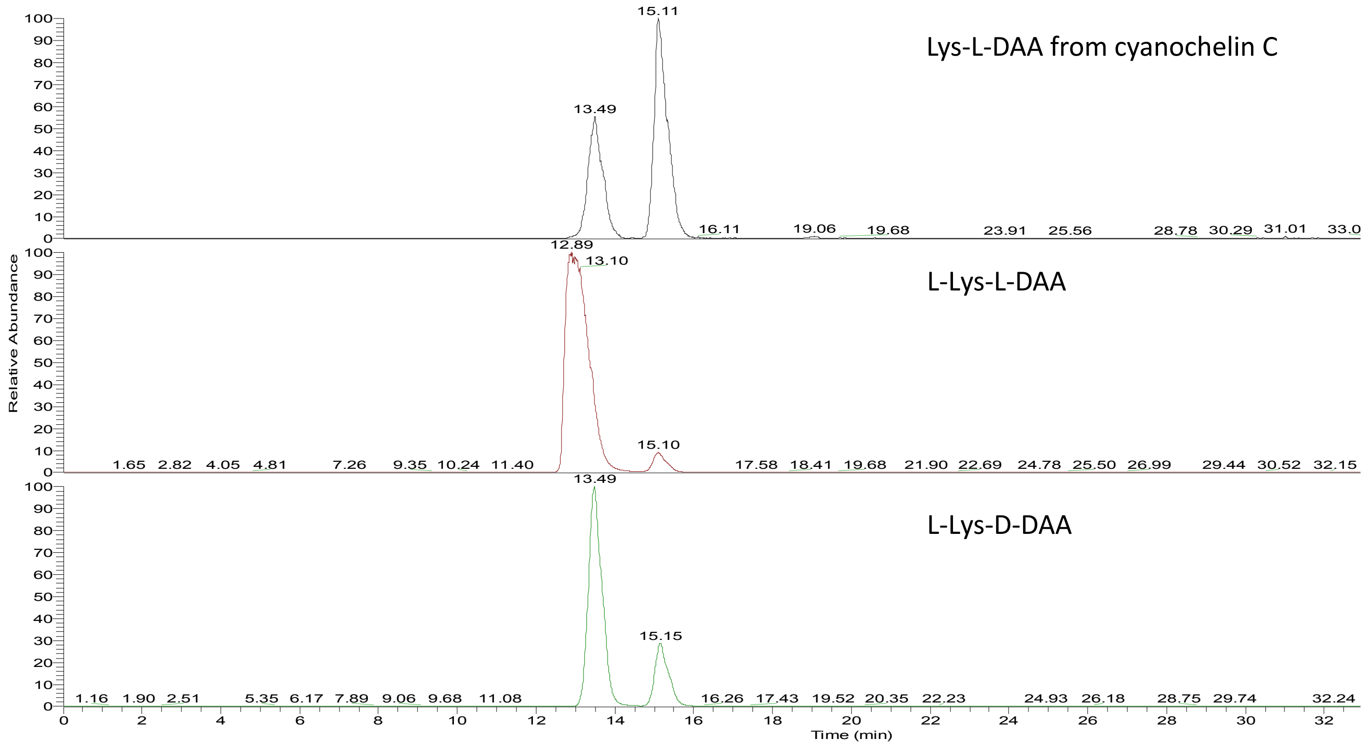
