## Supplementary figures and images for "Structural and Stereochemical Elucidation of Cyanochelin C, a Siderophore Associated with Novel Class of Cyanobacterial Acyl Hydrolases"

### Fig S11 - full phylogenetic tree of cyanobacterial acylases

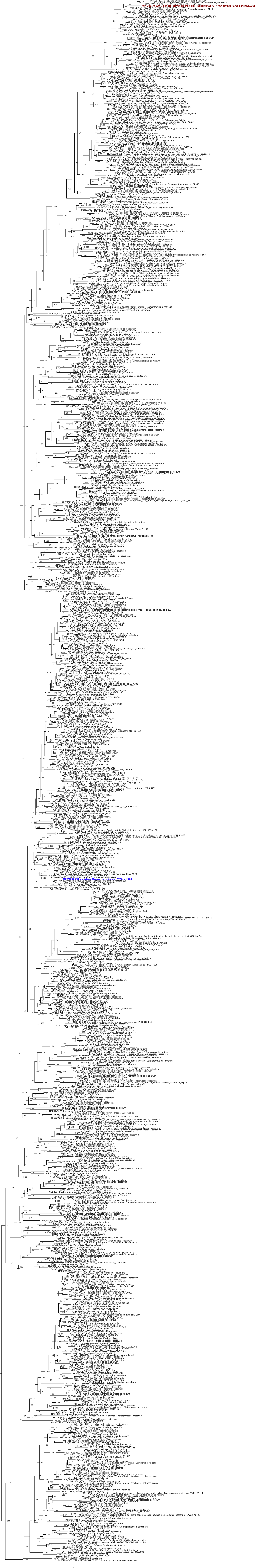
